# The LIF-LIFR Axis Promotes Liver Regeneration via Modulation of Angiogenesis and HGF Release from LSECs

**DOI:** 10.64898/2026.02.24.707802

**Authors:** Wenjing Zhou, Tanja Diemer, Hong Xin, Krishna Chaitanya Ginne, Kumar R.N. Naresh, Tommaso Mori, Nilima Biswas, Carlo Piermarocchi, Napoleone Ferrara

**Affiliations:** Department of Pathology, University of California, San Diego, La Jolla, CA 92093; Department of Ophthalmology, University of California San Diego, La Jolla, CA 92093; Department of Physics & Astronomy, Michigan State University, East Lansing, MI, USA – 48824

**Keywords:** Liver regeneration, LSECs, LIF, LIFR, HGF, VEGF, Angiogenesis

## Abstract

Liver sinusoidal endothelial cells (LSECs) play essential roles in liver regeneration after injury, but the underlying mechanisms remain incompletely defined. Here we report that leukemia inhibitory factor (LIF), which is rapidly induced after liver injury, acts as a key regulator of LSECs-driven liver regeneration through interaction with LSECs-enriched LIF receptor (LIFR). LIF directly stimulates LSECs proliferation and induces hepatocyte growth factor (HGF) release in a dose-dependent manner via LIFR signaling in LSECs, thereby indirectly promoting hepatocyte proliferation. Systemic LIF neutralization or endothelial cells (ECs)-specific *Lifr* loss impairs liver regeneration, whereas low-titer AAV-mediated LIF expression increases vascular density, elevates circulating HGF, and improves early liver recovery after partial hepatectomy (PHx) in mice. Together, these findings establish LIF-LIFR as a previously unrecognized endothelial axis to promote hepatocyte proliferation and suggest potential therapeutic strategies to enhance liver repair in patients.

**Highlights:**

1. LIF is upregulated after liver injury and LIF neutralization impairs liver recovery.
2. LIFR displays the highest expression in ECs; endothelial-specific *Lifr* deletion delays liver regeneration after injury.
3. LIF mediates a positive feedback loop including LSECs proliferation as well as HGF release via LIFR pathway.
4. LIF overexpression increases liver-to-body weight ratio in a dose-dependent manner and accelerates liver regeneration at early stage.

**Abstract Graphic:** 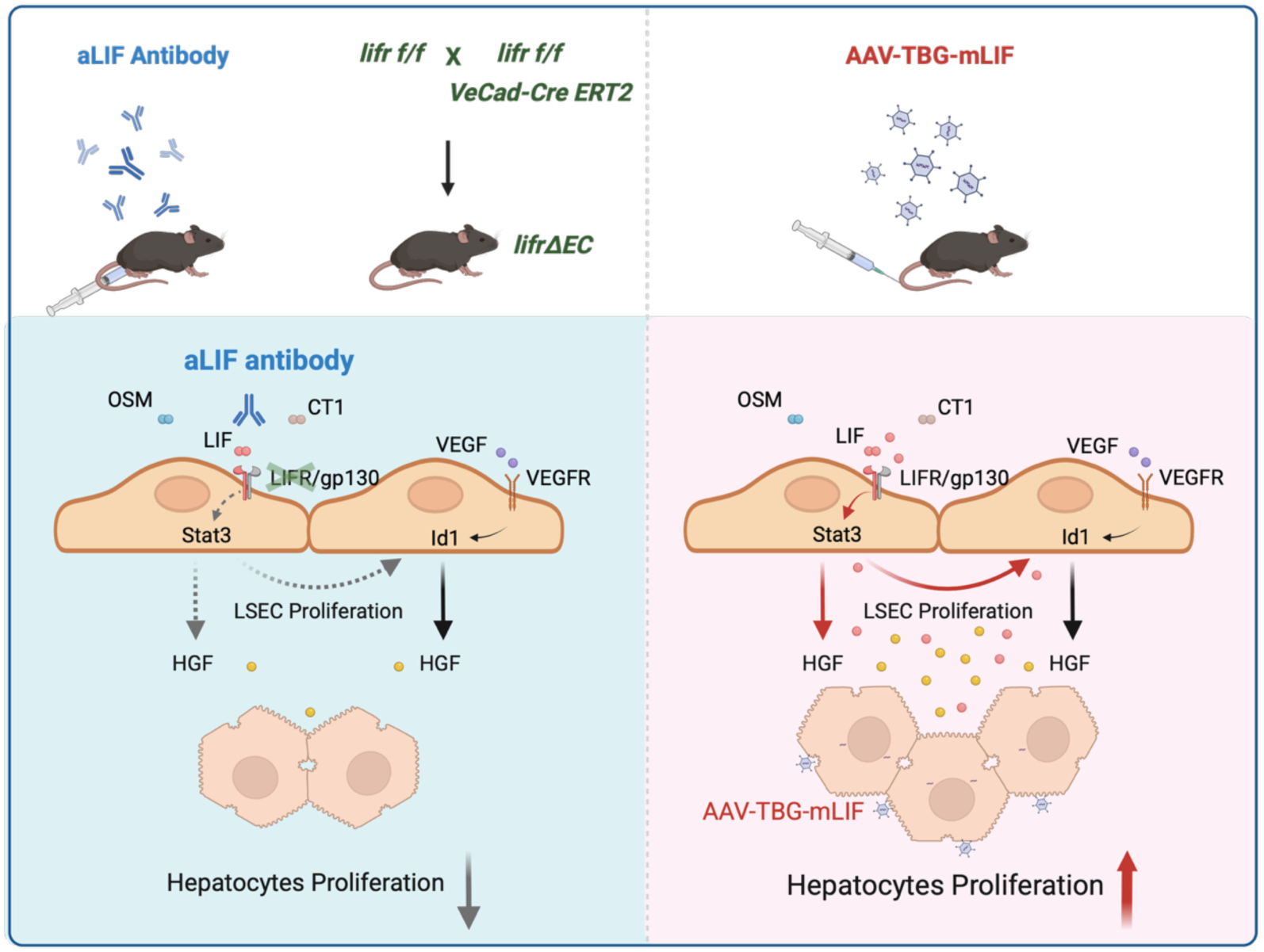

## Introduction

Unlike most organs, the liver has a remarkable regenerative capacity, which can lead to normal size restoration after surgical resection or acute injury (1, 2). Uncovering the mechanisms of liver regeneration is crucial for improving outcomes of patients with liver failure and for elucidating general regeneration principles, potentially applicable to aging and tissue engineering.

Liver ECs, as the most abundant component of non-parenchymal cells (NPCs), play essential roles in liver function and regeneration. Liver sinusoidal endothelial cells (LSECs) form a specialized, fenestrated, vascular network that spans periportal, mid-zonal and pericentral regions along the portal-central axis of the liver lobe and closely cooperate with both parenchymal and non-parenchymal cells to regulate hepatocyte proliferation (3–5). It is now well established that LSECs serve as an active signaling hub that coordinates both vascular remodeling and parenchymal regeneration through the release of paracrine/angiocrine cytokines (5–9).

Among LSEC-derived cytokines, hepatocyte growth factor (HGF) plays a major role as it directly stimulate hepatocyte proliferation (10). In the liver, HGF is produced mainly by NPCs, including hepatic stellate cells (HSCs), LSECs and Kupffer cells and is significantly upregulated following injury (11, 12). Additionally, conditional *hgf* depletion in LSECs results in impaired liver regeneration that cannot be compensated by HGF produced by other cell types (13, 14), indicating an essential LSEC-dependent contribution to the regenerative process.

Previous work has shown that vascular endothelial growth factor (VEGF) stimulates LSECs after injury. VEGF is mainly expressed in damaged and replicating hepatocytes, with additional contributions from hepatic stellate cells (HSCs), Kupffer cells and biliary epithelial cells (BECs), also known as cholangiocytes (15–17). VEGF binding to VEGFR2 in LSEC results in mitogenesis, angiogenesis and remodeling of the sinusoidal network (18, 19). Furthermore, in response to VEGF, LSECs release paracrine/angiocrine factors, notably HGF, that in turn stimulate hepatocytes proliferation, accelerating liver regeneration (9, 20–23). However, notwithstanding the importance of VEGF signaling, additional regulatory pathways governing LSEC activation during liver injury remain incompletely defined.

Leukemia inhibitory factor (LIF) is a cytokine that belongs to the interleukin-6 family and has complex and context-dependent effects on stem cell homeostasis, regeneration, immunity and cancer development (24). The functions vary in different tissues and cells, and even within the same cell type. For example, early studies characterized LIF as a growth inhibitor for aortic ECs (25), but more recent studies have reported unexpected mitogenic effects on certain ECs types such as choroidal ECs (26).

LIF specifically interacts with the LIF receptor (LIFR), triggering the formation of a heterodimeric complex composed of gp130 and LIFR, which activates JAK/STAT3, ERK and PI3K pathways to support cell survival and proliferation (27, 28). *lifr/LIFR* expression is relatively high in the liver among different organs (29). Moreover, analysis of Tabula Muris indicates that *lifr* is particularly enriched in LSEC among ECs from multiple mouse organs (26). However, studies on LIF-LIFR as well as other IL-6 family cytokines in the liver have largely focused on hepatocyte responses (30, 31), rather than on LSEC, leaving it a comparatively underexplored area during liver injury.

We describe here LIF as a previously unrecognized mitogen for LSECs that also stimulates HGF release from LSECs, specifically via LIFR-STAT3 signaling. This VEGF independent LIF/LIFR circuit is essential for hepatocytes proliferation during liver injury, providing new insights into endothelial-parenchymal crosstalk and suggesting a potential therapeutic approach.

## Results

### *lif/LIF* is induced in mice and humans after liver injury

To explore whether LIF signaling may be implicated in liver recovery, we analyzed a mouse single cell RNA-seq dataset (GSE223558) with multiple time points following acetaminophen (APAP)-induced liver injury. We found that *lif* expression is very low at basal levels but starts to increase ∼18 hours post-APAP and returns to baseline by ∼48 hours (Figure 1A). A similar increase in *lif* was also observed in the CCl_4_-induced liver injury models. In addition to *lif*, other IL-6 family cytokines such as *il6*, *osm* and *cntf* display slight increases following CCl_4_-induced liver injury in mice (GSE233084) (Figure S1A). Further analysis across different cell types in injured liver indicates that *lif* upregulation occurs in multiple cell populations, including cholangiocytes and hepatocytes (Figure 1B), while *vegfa* is mainly expressed in hepatocytes, in both healthy and injured mouse livers (Figure S1B).

**Figure 1.**
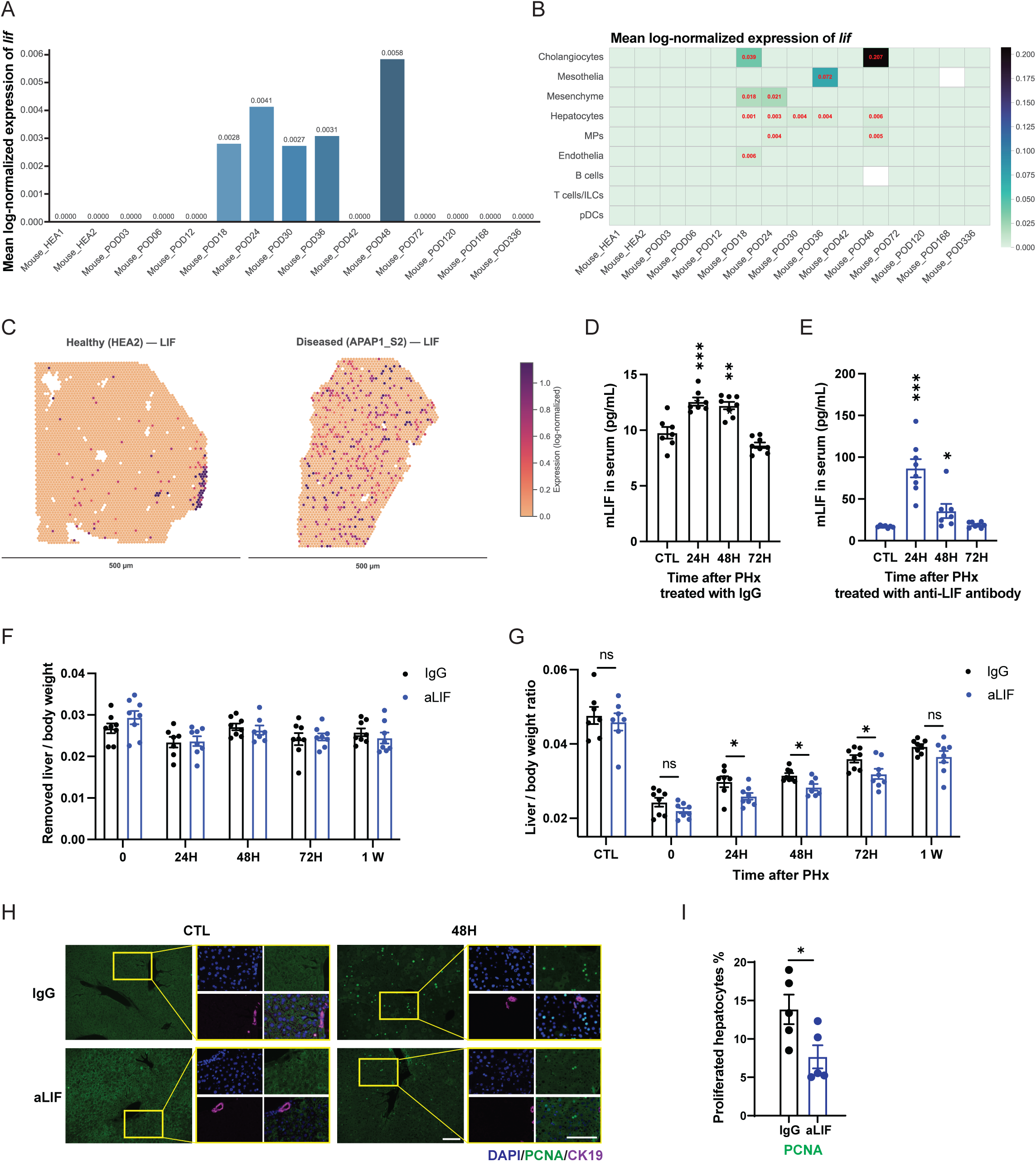
Injury-induced LIF is required for liver regeneration. (A) *lif* expression in the scRNA-seq from mouse livers with acetaminophen (APAP)-induced injury at the indicated time points. (B) *lif* expression in the scRNA-seq across cell types in the livers of mouse with APAP injury at the indicated time points. (C) *lif* expression in spatial transcriptomics of liver samples from healthy (HEA2) and APAP injury (APAP1_S2) humans. (D&E) Serum mLIF levels at 24, 48, and 72 h after partial hepatectomy (PHx) in mice treated with IgG (D) or anti-LIF neutralizing antibody (E). (F) Resected liver-to-body weight ratio in mice treated with anti-LIF antibody or control IgG. (G) Liver-to-body weight ratio at 24, 48, 72 h and 1-week post-PHx in mice treated with anti-LIF antibody or control IgG. (H) Immunofluorescence staining of liver sections at 48 h post-PHx from mice treated with anti-LIF antibody or control IgG. DAPI (blue), PCNA (red), CK19 (purple). Scale bars: 100 μm. (I) Quantification of PCNA-positive hepatocytes ratio at 48 h post-PHx in mice treated with anti-LIF antibody or control IgG.

Consistent with these findings in mice, analysis of a human liver dataset (GSE223581) revealed increased *LIF* expression after APAP-induced injury (Figure S1C). Cell type analysis within the liver further showed that the *LIF* upregulation was mainly contributed by cholangiocyte (Figure S1D). In contrast, *VEGF* upregulation in injured human livers was mainly driven by hepatocytes (Figure S1E). Spatial Transcriptomics analysis of a human dataset (GSE223559) independently confirmed low basal *LIF* levels, with marked increases after APAP-induced injury (Figure 1C), and cell type analysis again identified cholangiocytes as the dominant source of *LIF* in both healthy and injured human livers (Figure S1F).

We next validated injury-associated LIF induction in a mouse liver regeneration model. Following partial hepatectomy (PHx), serum LIF levels detected by ELISA increased rapidly within 24 hours post-PHx and returned to baseline by 72 hours (Figure 1D). Together, these data demonstrate that LIF is an injury inducible cytokine in both mice and humans with liver damage.

### LIF neutralization impairs liver regeneration after PHx

To determine whether injury-associated LIF elevation was functionally required during liver injury, we administered the anti-LIF neutralizing monoclonal antibody (Mab) D25 to mice before PHx. Mab D25, originally described by Genentech, Inc. (32), was purified in our lab from hybridoma cell supernatants obtained from the ATCC (see Methods), and key findings were verified using purified Mab D25 provided by Genentech. Due to stabilization after binding with the antibody, detectable serum LIF levels were higher in anti-LIF treated mice compared to IgG control groups and displayed more obvious increases at 24- and 48-hours post-injury (Figure 1E). The specificity of the inhibitory effects of Mab D25 was confirmed in subsequent assays (Figure S3B-S3D).

Following short-term systemic anti-LIF antibody treatment, major organs, body weight as well as liver-to-body weight ratio didn’t show obvious changes (CTL in figure 1G). Liver histology by H&E staining was similar in control and treated groups (Figure S1I). After PHx, anti-LIF treatment reduced liver-to-body weight ratio at 24, 48, 72 hours and 1-week compared with IgG controls (Figure 1G), starting with a comparable removed liver-to-body weight ratio (Figure 1F), indicating impaired liver regeneration after injury. Similar effects were observed with Mab D25 obtained from Genentech. (Figure S1G&S1H). Consistent with impaired regeneration, serum AST levels increased at 24 hours post-PHx upon LIF neutralization compared with IgG controls (Figure S1J). Immunofluorescence staining for PCNA in liver sections further revealed reduced hepatocyte proliferation at 48 hours in mice treated with anti-LIF antibody (Figure 1H, quantified in Figure 1I).

### *Lifr/LIFR* is enriched in liver ECs in mice and humans

We next examined the expression pattern of LIF receptor within liver. We analyzed *lifr* expression across different cell types in the whole liver from mouse single cell RNA-sequencing (scRNA-seq) data (GSE223558) and found that *lifr* expression was most enriched in ECs, followed by dendritic cells, mesenchyme and hepatocytes (Figure 2A, upper). The co-receptor *il6st (gp130)* also showed the highest expression in ECs (Figure 2A, lower). In agreement with previous studies, *kdr* and *flt* displayed the highest expression in endothelia, while *il6ra* was primarily observed in hepatocytes (Figure 2A, lower). The specific *lifr* enrichment in ECs in the liver was independently confirmed with another mouse single-cell dataset (GSE151309) (Figure S2A).

**Figure 2.**
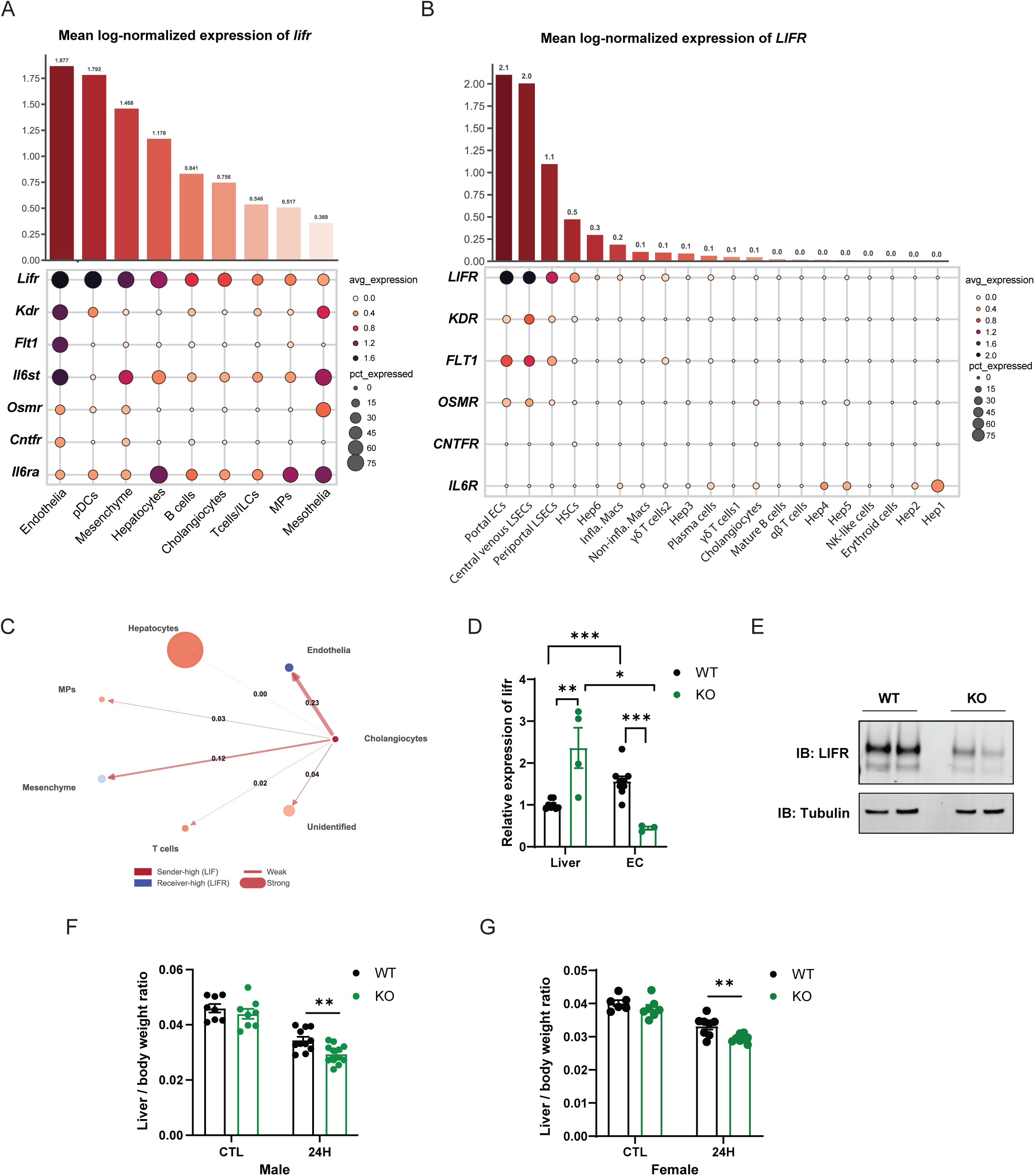
LIFR displays the highest expression in ECs in the liver and endothelial LIFR deletion impairs regeneration. (A) Expression of IL-6 family receptors in scRNA-seq across cell types of livers from healthy (HEA1+HEA2) mice. The top panel is a bar plot of *lifr* expression, the bottom panel is a bubble plot of multiple receptors as indicated. (B) Expression of IL-6 family receptors in scRNA-seq across cell types of livers from healthy human. The top panel is a bar plot of *LIFR* expression, the bottom panel is a bubble plot of multiple receptors as indicated. (C) Circular network plot of *LIF-LIFR* interaction between different cell types in liver from healthy human (HEA5). (D) qPCR of *lifr* in isolated LSECs and whole liver from *lifr f/f* and *lifrΔEC* mice. (E) Western blot of LIFR in isolated LSECs from *lifr f/f* and *lifrΔEC* mice. (F) Liver-to-body weight ratio at 24 h post-PHx in male *lifr f/f* and *lifrΔEC* mice. (G) Liver-to-body weight ratio at 24 h post-PHx in female *lifr f/f* and *lifrΔEC* mice.

In a human liver scRNA-seq (GSE115469), *LIFR* exhibited the highest expression in ECs, including periportal LSEC (Zone1), central venous LSEC (Zone 2/3) and portal ECs, followed by lower expression in hepatic stellate cells (HSCs) (Figure 2B, upper). As in mouse liver, the canonical VEGF receptors *KDR* and *FLT1* expression was largely restricted to ECs, whereas *IL6R* expression was mainly in hepatocytes (Figure 2B, lower), highlighting diverse functions of IL-6 family receptors with a compartmentalized distribution. The distinct localization was independently confirmed in an additional human liver scRNA-seq dataset (GSE223581) with *LIFR* enrichment in EC (Figure S2B). Cell-cell communication analysis of healthy human livers (GSE223581) indicated that the predicted *LIF* - *LIFR* interaction was most prominent between cholangiocytes and ECs (Figure 2C & S2C-S2E).

Collectively, these data point to EC as the principal *lifr/LIFR* expression compartment in both mice and humans, and support the hypothesis of intense communication between cholangiocytes and ECs, indicating the LIFR signaling axis functions as an endothelial centered pathway, different from the hepatocytes focused IL6R signaling.

### Loss of endothelial *lifr* impairs liver regeneration after PHx

To further characterize the LIFR responses during liver injury, we first monitored *lifr* expression levels following PHx. In the whole liver, *lifr* decreased transiently at 24-48 hours and recovered by 72-hours after PHx (Figure S2H, left). Because LSECs have the highest LIFR expression, we isolated primary LSECs from liver after PHx at indicated time points. The purity of LSECs was validated by qPCR and FACS using specific markers (Figure S2F&S2G). *Lifr* expression in isolated LSECs displayed similar trends as in whole liver after PHx (Figure S2H, right), indicating LSECs were involved in LIFR response upon liver injury.

Based on the data above, we hypothesized that LSECs may be involved in liver recovery through LIF-LIFR signaling. To test this hypothesis, we generated ECs-specific *lifr* knockout mice (*lifrΔEC*) by crossing *lifr f/f* mice with tamoxifen inducible *VeCad-CreERT2* mice and inducing Cre expression at 8 weeks old. qPCR analysis of liver and isolated LSECs from *lifr f/f* (indicated as WT) and *lifrΔEC* (indicated as KO) showed the strong enrichment of *lifr* in isolated LSECs compared with whole livers in *lifr f/f* mice, and a significant reduction of *lifr* in LSECs from *lifrΔEC* mice. Additionally, in whole liver, the *lifr* expression increased in *lifrΔEC* mice compared with *lifr f/f*, indicating compensatory expression of *lifr* from other cell types (Figure 2D). Reduced LIFR protein in LSECs from *lifrΔEC* was validated by western blot (Figure 2E).

In *lifrΔEC* mice, the baseline liver-to-body weight ratio didn’t change between genotypes (CTL from Figure 2F and Figure 2G). After PHx, the liver-to-body weight ratio was significantly reduced at 24-hour post-surgery in both male and female *lifrΔEC* mice (Figure 2F & 2G), indicating that endothelial LIFR is required for liver regeneration. Collectively, these data support the hypothesis that the LIF-LIFR axis is involved in liver recovery after injury. Additionally, LSECs, which have the highest LIFR expression, serve as an essential target of LIF to coordinate cell-cell response in the liver regenerative process.

### LIF is a mitogen for LSEC but does not directly stimulate hepatocyte proliferation

To explore the mechanisms of LIF-LIFR signaling in LSECs, we treated primary LSECs with recombinant LIF. LIF promoted LSECs proliferation in a biphasic (bell-shaped) dose-response, with effects starting around 0.3 ng/mL, reaching a plateau at around 10 ng/mL and declining at higher concentrations (Figure 3A). LIF-induced proliferative effects were blocked by Mab D25 (Figure S3B&S3D), which did not show any inhibitory effects on VEGF-induced proliferation (Figure S3C). Similarly, the VEGF-neutralizing antibody B.20.4.1 showed clear inhibition of VEGF induced LSECs proliferation, without inhibition of LIF effects (Figure S3E&S3F). Additionally, other IL-6 family cytokines, which also interact with LIFR, including OSM and CT-1, had similar effects as LIF in stimulating LSEC proliferation (Figure 3B&3C). IL-6 displayed only modest proliferative effects at high concentrations, weaker than LIFR-engaging ligands (Figure S3A). Genetic loss of *lifr* in LSECs impaired proliferation in response to LIF (Figure 3D), without affecting VEGF induced proliferation (Figure 3E&3F).

**Figure 3.**
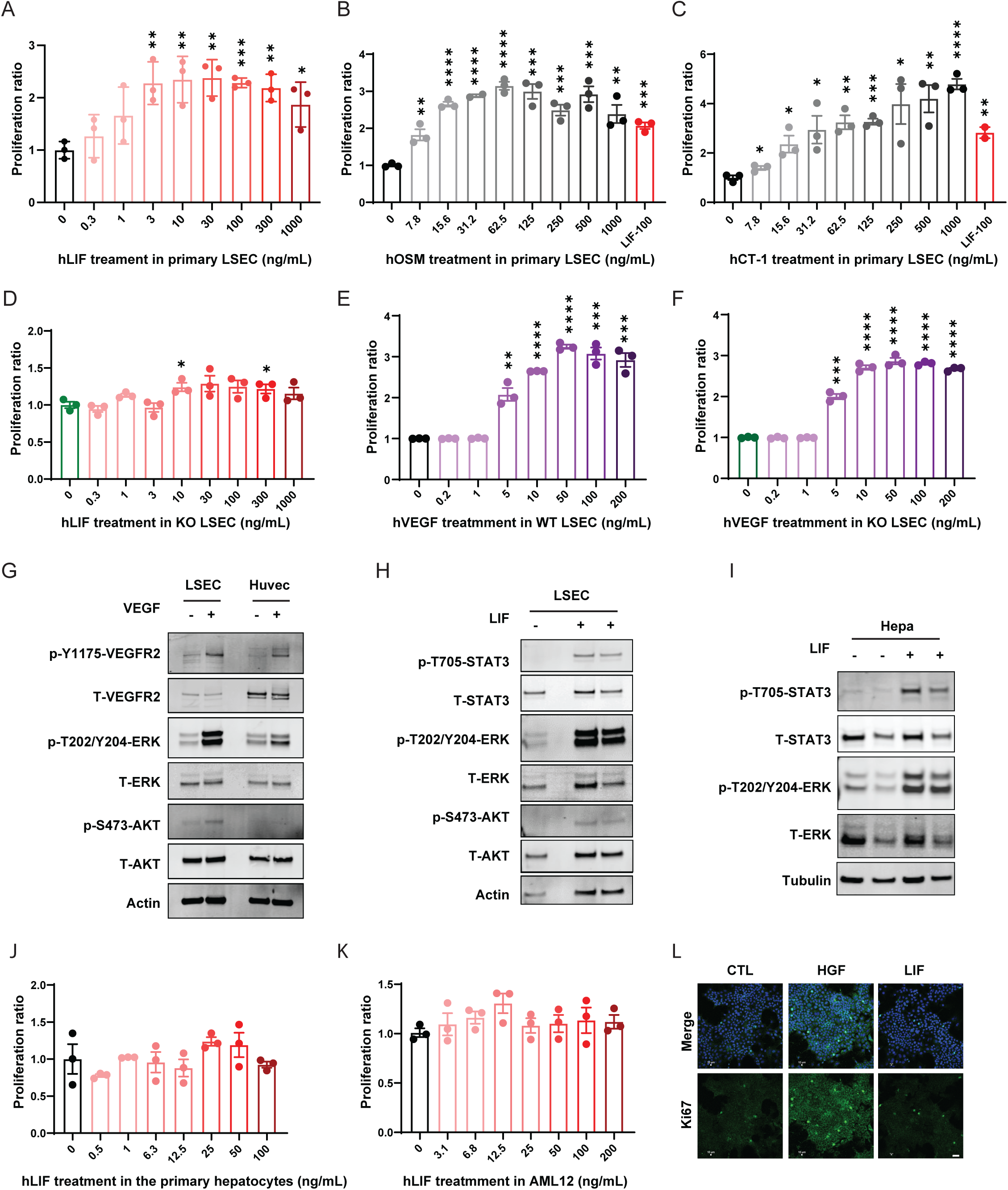
LIF directly promotes LSEC proliferation via LIFR but does not directly promote hepatocytes proliferation. (A-C) Proliferation assays of primary LSECs treated with increasing doses of hLIF (A), hOSM (B) or hCT-1 (C). 100 ng/mL hLIF was used as positive control. (D) Proliferation assays of LSECs isolated from *lifrΔEC* mice treated with increasing doses of hLIF. (E&F) Proliferation assays of LSECs isolated from *lifr f/f* (E) or *lifrΔEC* (F) mice treated with increasing doses of hVEGF. (G&H) Western blot analysis of primary LSECs treated with hVEGF (G) or hLIF (H) (I) Western blot of primary hepatocytes treated with hLIF. (J) Proliferation assays of primary hepatocytes treated with increasing doses of hLIF. (K) Proliferation assays of AML12 cells treated with increasing doses of hLIF. (L) Immunofluorescence staining of AML12 cells treated with 80 ng/mL hHGF or hLIF. DAPI (blue), Ki67 (green). Scale bar: 50 μm.

VEGF-induced VEGFR2 tyrosine phosphorylation and activation of downstream proliferative pathways in LSECs have been previously characterized (9, 33). Western blot analysis confirmed that isolated primary LSECs exhibit a typical phosphorylation response to VEGF, comparable to that observed in HUVECs (Figure 3G). LIF treatment (40 ng/mL) also induced phosphorylation of STAT3, ERK and AKT in primary LSECs (Figure 3H). In primary hepatocytes, LIF likewise induced STAT3 and ERK phosphorylation (Figure 3I), raising the possibility of direct hepatocyte responses. However, in isolated primary hepatocytes in which proliferation was induced by the addition of HGF and EGF, LIF had no significant proliferative effects (Figure 3J&S3G). Similarly, in the AML12 mouse hepatocyte cell line, LIF failed to stimulate proliferation (Figure 3K) and Ki67 induction (Figure 3L) at concentrations in which HGF significantly stimulated proliferation (Figure S3H).

Together, these data indicate that, alongside VEGF, LIF is a potent mitogen for LSECs via activation of LIFR dependent signaling but does not directly drive hepatocyte proliferation. The bell-shaped dose-response to LIF has been described in previous reports (34) and is commonly observed among other IL6 family cytokines (35, 36), underscoring the need for precise dosing of these signals.

### LIF promotes hepatocyte proliferation indirectly by paracrine induction of HGF from LSECs via LIFR-STAT3 Pathway

LSECs serve as a key communication hub with other liver cell types via secretion of multiple cytokines during injury. To elucidate the mechanisms underlying LIF effects on cell-cell crosstalk via cytokines, secretome analysis was performed in primary LSECs, treated or not with 40 ng/mL hLIF. Principal Component Analysis (PCA) revealed clear separation between the two groups (Figure S4A). Differential analysis revealed 13 upregulated and 12 downregulated proteins shown in volcano plot (Figure 4A). Among these, HGF was significantly increased.

**Figure 4.**
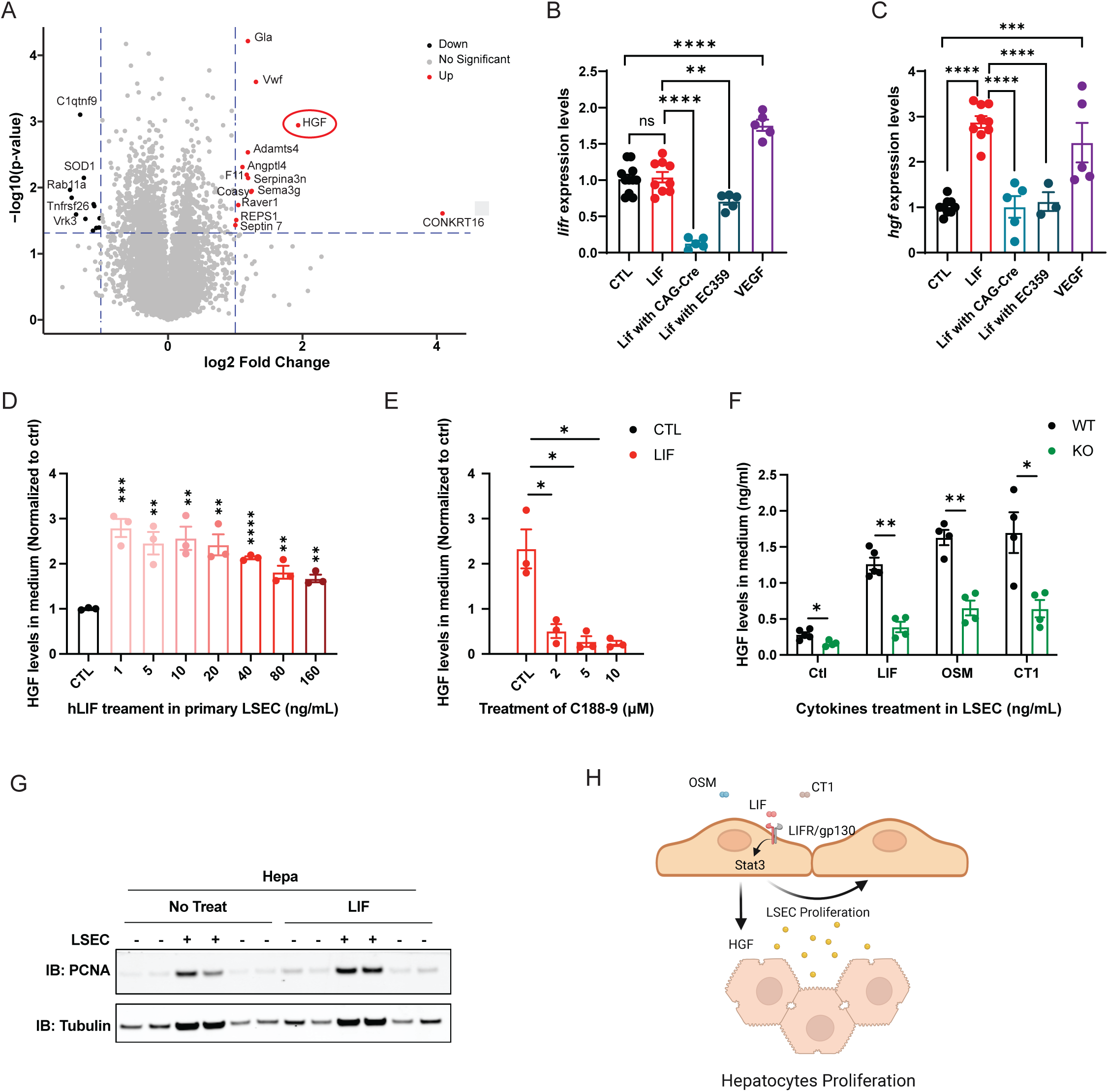
LIF induces hepatocyte proliferation indirectly via paracrine release of HGF from LSECs. (A) Volcano plot of differentially secreted proteins from primary LSECs treated with 40 ng/mL LIF versus control (n = 4). (B&C) qPCR of *lifr* (B) and *hgf* (C) levels in LSECs isolated from *lifr f/f* mice with indicated treatments. (D) HGF concentrations in conditioned medium from LSECs treated with hLIF at the indicated doses. (E) HGF concentrations in conditioned medium from LSECs treated with 40 ng/mL hLIF in the presence of the STAT3 inhibitor C188-9 at the indicated doses. (F) HGF concentrations in conditioned medium from LSECs isolated from *lifr f/f* or *lifrΔEC* mice treated with hLIF, hCT-1 or hOSM. (G) Western blot analysis of hepatocytes co-cultured with LSECs with or without 40 ng/mL LIF. Quantification is shown in Figure S4C. (H) Model depicting LIF action on LSECs to stimulate angiogenesis and HGF release, thereby supporting hepatocyte proliferation via paracrine regulation.

Consistent with proteomic results, LIF increased *hgf* mRNA levels in LSECs, comparable to that induced by VEGF (Figure 4C). Analysis of medium treated with different LIF concentrations showed that HGF release followed a biphasic (bell-shaped) dose-response, with maximal induction at low concentrations and relatively reduced release at higher concentrations (Figure 4D), consistent with the noted LIF effects on proliferation. LIF-induced HGF upregulation was suppressed by disrupting *lifr* function,either by *Cre*-mediated recombination (AAV-CAG-Cre) or using a small molecule inhibitor (EC359) (Figure 4B&4C). Moreover, blockade of STAT3 signaling with C188-9 prevented LIF-induced HGF release (Figure 4E), supporting a LIFR-STAT3 dependent mechanism. Interestingly, even without the addition of LIF, baseline HGF levels were decreased in LSECs isolated from *lifrΔEC* mice compared with *lifr f/f* controls (Figure 4F&S4B), suggesting autocrine mechanisms contributing to HGF release via the LIFR pathway. OSM and CT1 also showed a strong induction of HGF expression in LSECs, even higher than LIF at the same dose, and this induction was similarly LIFR-dependent (Figure 4F&S4B).

To determine the role of LSECs-secreted HGF on hepatocyte proliferation, primary LSECs were co-cultured with primary hepatocytes. Co-culture significantly promoted hepatocyte proliferation and LIF addition further enhanced hepatocyte proliferation as shown by higher PCNA expression (Figure 4G, quantified in S4C), supporting an indirect mechanism of LIF on hepatocyte proliferation via HGF signaling. Additionally, mHGF or hHGF treatment did not stimulate primary LSECs proliferation (Figure S4D & S4E), indicating that LIF induced LSECs proliferation is not mediated by HGF.

Together, these data demonstrated the dose-dependent role of LIF in LSECs angiogenesis and paracrine HGF secretion via activation of LIFR-STAT3 pathway, thereby coupling vascular expansion and hepatocytes proliferation (Figure 4H).

### Low-dose LIF expression accelerates liver regeneration at early stage

To test the therapeutic potential and dose-dependence of LIF actions in vivo, we used AAV8-TBG-mLIF to drive mLIF expression in hepatic cells at different titers. At lower titers, mice maintained normal body weight and activity (Figure 5A). Interestingly, low-titer LIF increased liver weight as well as liver-to-body weight ratio in a dose-dependent manner (Figure 5B-5C) which is in line with increased levels of LIF in serum (Figure 5D). Liver histology and serum AST levels did not show any obvious abnormalities in response to systemic low-dose LIF expression (Figure S5E & CTL in S5G). Spleens also showed similar dose-dependent increases as the liver (Figure S5A & S5B), whereas kidneys were unchanged (Figure S5C&S5D). However, higher LIF expression (AAV titers over 1×10^9 GC/mice) led to reduced body, liver, spleen and kidney weights and signs of poor health (Figure 5A-5B &S5A-S5D). Low-dose LIF induced increase of liver to body weight ratio was no longer observed at higher doses (Figure 5C), consistent with the content-dependent response to LIF.

**Figure 5.**
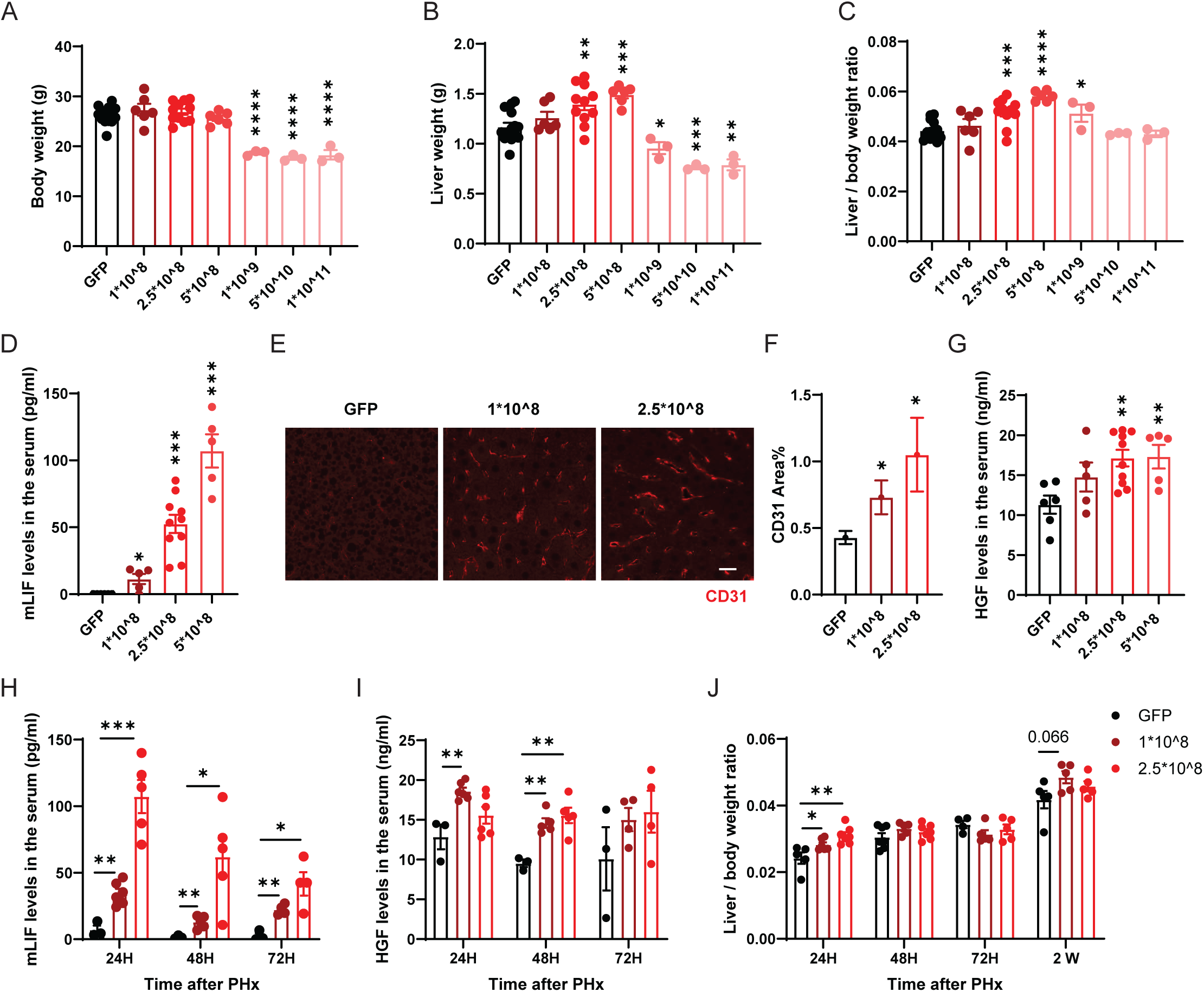
Low-dose LIF expression accelerates liver regeneration at early stage. (A-C) Body weight (A), liver weight (B) and liver-to-body weight ratio (C) of mice injected with AAV8-TBG-mLIF at the indicated titers. (D) Serum mLIF levels in mice injected with AAV8-TBG-mLIF at the indicated titers. (E&F) CD31 immunofluorescence staining of liver sections from mice injected with low-doses AAV8-TBG-mLIF (1×10⁸ or 2.5×10⁸ GC/mouse) (E) and quantified in (F). Scale bar=50 μm. (G) Serum HGF levels in mice injected with AAV8-TBG-mLIF at the indicated titers. (H) Liver-to-body weight ratio at the indicated time points post-PHx in mice injected with low-doses AAV8-TBG-mLIF (1×10⁸ or 2.5×10⁸ GC/mouse). (I&J) Serum mLIF (H) and HGF (I) levels at 24, 48 and 72 h post-PHx in mice injected with low-doses AAV8-TBG-mLIF (1×10⁸ or 2.5×10⁸ GC/mouse).

Serum mLIF levels increased with AAV titers, reaching around 10-fold with 1×10^8 GC/mice, 50-fold with 2.5×10^8 GC/mice and 100-fold with 5×10^8 GC/mice over baseline (Figure 5D). Consistent with LIF’s mitogenic effects on LSECs, CD31 immunofluorescence demonstrated increased vascular density with rising LIF levels (Figure 5E, quantified in 5F). Serum HGF levels also increased with LIF dosage but approached a plateau at the highest tested titers (Figure 5G).

To further test the protective potential of LIF against liver injury, we performed PHx in mice with LIF expression at lower titers. Consistent with the data shown in Figure 1D&E, serum LIF levels peaked 24 hours post-surgery at different titers of LIF as indicated (Figure 5H). Serum AST levels were improved at different time points (Figure S5G), showing better recovery of liver function upon LIF elevation. Notably, serum mHGF levels increased according to the LIF levels, consistent with LIF effects on HGF release in vitro (Figure 5I). The liver to body weight ratio was elevated at 24 hours post-PHx with LIF expression at different titers (Figure 5J), including after normalization to pre-surgery differences (Figure S5F). However, at 48 and 72 hours after PHx, LIF overexpression showed a trend to impaired liver regeneration in normalization analysis, most evident at higher titer (Figure S5F). At 2-week post-PHx, LIF overexpression increased liver to body weight ratio, in which the lower titer was better than that of higher titer (Figure 5J&S5F).

In summary, we conclude that LIF overexpression improves liver recovery in a dose- and time-dependent manner, with a low dose window that maximizes vascular expansion, HGF induction and early liver growth.

## Discussion

Liver regeneration is orchestrated by dynamic crosstalk among hepatocytes and their surrounding microenvironment (37). In the liver, LSECs are central mediators of regeneration, not only by responding to cytokines released from LSECs themselves or other cell types such as hepatocytes or cholangiocytes, but also by releasing angiocrine factors such as HGF, Ang2, Wnt9b, Wnt2, and Notch ligands (6, 9, 38–40). Among the angiocrine signals, HGF plays a pivotal role as hepatocyte mitogen (41), while VEGF-VEGFR signaling has been regarded as a key regulator of LSECs activation and HGF release after liver injury (9). Here, we integrate and extend established models of LSECs-mediated regeneration by identifying a VEGF-independent LIF-LIFR centered pathway that induces dose-dependent endothelial expansion and paracrine HGF release. This raises the possibility that the LIF and VEGF signaling pathways may act in parallel, and that their co-activation could produce synergistic or dose-dependent effects that shape the efficacy and timing of regeneration.

Our study underscores the importance of dynamic cell-cell communication during liver regeneration. Transcriptomic and cell-cell interaction analyses suggest that injury-induced LIF is likely produced by cholangiocytes and interact with LIFR enriched LSECs. As an important ligand source, cholangiocytes have been implicated in influencing the liver injury microenvironment by promoting immune cell infiltration and HSCs activation (42). Cholangiocytes also engage in crosstalk with ECs, involving, for example, endothelin-1 (ET-1) signaling (43) and VEGF-dependent autocrine or paracrine circuits that support biliary expansion and vascular adaptation (16). Although the major cellular source of LIF after liver injury remains to be further defined using cell type specific analysis or lineage tracing, our findings suggest that cholangiocytes may communicate with LSECs through multiple complementary signals including LIF.

More broadly, our findings highlight LSECs as an active hub for IL6 family cytokines in liver regeneration, whereas most previous studies focused on hepatocyte responses (31, 44). The distinct cellular localization of LIFR is enriched in LSECs, while IL6R is mainly expressed in hepatocytes, supporting the divergent and cell type specific functions during liver repair. In addition to LIF, we found that other LIFR-engaging cytokines, including CT-1 and OSM (45), elicit effects similar to LIF in LSECs, indicating that LIFR may act as a broader endothelial receptor integrating multiple IL-6 family ligands during injury.

Notably, LSECs exhibit bell-shaped, dose-dependent responses to LIF, which may be helpful for the tight coordination between vascular remodeling and hepatocyte proliferation while preventing excessive signaling during regeneration. Such biphasic regulation possibly reflects the need for intermediate ligand levels to maximize downstream outputs, whereas higher levels induce intrinsic brakes such as negative signaling feedback and receptor desensitization to dampen signaling and limit pathological overactivation(46–48). Similar biphasic dynamics have been described for LIF (26, 34), other IL6 family cytokines (36, 49), as well as VEGF (46), underscoring the importance of dose in translating cytokine signals into effective outcomes with safety.

At the signaling level, LIFR engages gp130 to classically activate the JAK-STAT3, ERK/MAPK and AKT-mTOR pathways. In primary LSECs, LIF triggered rapid and robust STAT3 and ERK phosphorylation, suggesting LIF as a potent cytokine that can directly activate regenerative signaling within endothelium. Importantly, STAT3 inhibition further supports the direct HGF release induced by LIF in LSECs. This is consistent with previous work in HEK293 cells, which showed that STAT3 induction directly activates the *hgf* promoter and increases HGF expression (50). Our findings establish a direct link between LSECs centered LIF-LIFR signaling and paracrine HGF release, thereby promoting hepatocyte proliferation. Moreover, the STAT3 dependent response, unlike Id1 activation, supports the VEGF-independent mechanism for inducing HGF in LSECs.

An apparently puzzling observation is the lack of a proliferative response to LIF in hepatocytes despite strong STAT3 phosphorylation and downstream signaling events. However, recent studies report that *lifr* deficiency or overexpression in hepatocytes does not directly affect hepatocyte proliferation response to HGF ex vivo or in vitro (31). Other studies also show that IL6 alone is not able to stimulate hepatocyte growth (51, 52). Interestingly, LIF-induced STAT3 activation induced proliferation of BCE via the LIFR-JAK/STAT3 pathway, whereas activation of the same pathway in BAE cells led to growth inhibition with distinct gene expression pattern (26), further indicating that LIF-induced STAT3 phosphorylation may result in markedly different effects in different cell types.

Our in vivo gain of function data emphasize the tight dose- and time-dependent roles of LIF in regeneration. High titers of LIF expression in mice result in weakness and significant reductions in body and organ weight, consistent with the LIF-induced cachexia phenotype reported previously (53). In contrast, our findings with low dose LIF expression, show no obvious systemic toxicity but instead increased vascular density and liver-to-body weight ratio in a content-dependent manner and improved early functional recovery after PHx. At later stages, however, higher LIF doses delayed regeneration, supporting the physiological relevance of biphasic responses we observed in vitro. This is consistent with prior reports that excessive IL6/STAT3 pathway inhibit liver regeneration (54). Additionally, endogenous LIF is a transient, rapidly turnover signal and it returns to basal levels as regeneration progresses, whereas AAV mediated expression led to a sustained LIF exposure which may extend beyond the physiological window. This kinetic mismatch provides an additional explanation for the suppression of liver repair at later stage. LIF also led to dose-dependent spleen enlargement, as reported previously (55), which could secondarily modulate hepatic immune environment and thereby influence regeneration (56). The mechanisms of LIF-induced liver growth are complex and likely involve multicellular interactions. Testing LIF gain-of-function in *lifrΔEC* mice will help to quantify the contribution from LSECs.

In summary, our work defines the endothelial LIF-LIFR axis as a central, dose-and time-sensitive regulator of liver regeneration, forming a positive feedback loop between LSEC expansion and hepatocyte proliferation via paracrine HGF release. These findings offer a rationale for developing controlled strategies to modulate endothelial LIF-LIFR signaling to enhance liver repair in clinical settings where effective regenerative therapies remain limited.

## Methods

### Animals

Animal studies were approved and performed in compliance with guidelines of the University of California San Diego (UCSD) Institutional Animal Care and Use Committee (IACUC). Male C57BL/6J mice (8 weeks old) were purchased from the Jackson Laboratory (Bar Harbor, ME, USA). *Lifr f/f* mice (gift from Kyunghee Choi; generated as described previously (57) and VE-Cadherin-Cre mice (gift from Miguel Lopez-Ramirez at UCSD; generated by Ralf H. Adams) were maintained on a C57Bl/6J background. Genomic DNA from ear punches or tail was used for genotyping *Lifr f/f, VE-Cadherin-Cre* mice. PCR was carried out using 2 x Rapid Taq Master Mix (Vazyme, P222-02). Primers for identification of Cre positive offspring VE-Cadherin-Cre F 5’-CTGCATTACCGGTCGATGCA-3’ and VE-Cadherin-Cre R 5’-ACGTTCACCGGCATCAACGT-3’ produces a 350 pb PCR fragment. *Lifr f/f* genotyping was carried out as described. Briefly, primer *Lifr* CKO up-forward (U5) 5’-CCTTTTGCAGTCTCACAGGTCTTG-3’, *Lifr* CKO-up reverse (U3) 5’-TGGAAAGCAGACAAGCACAAGC-3’, *Lifr* CKO-down reverse (D3) 5’-TGGTAAGTCCAACTCAGCTAA GTG C-3’ R were used in two separate genotyping PCR reactions to detect wild type (368 bp) or *loxP* (461 bp) containing bands with (U3) and (U5) primers and primer pair (U5) and (D3) to detect the null *Lifr* Δ/Δ allele (474 bp). Tamoxifen (Millipore-Sigma, T5648), dissolved at 20 mg/mL in corn oil, (Millipore-Sigma, C8267) was used to induce the endothelial specific *LifR* knock-out. Mice received 0.225 mg Tamoxifen/g each day by gavage for 5 consecutive days. Experiments were carried out 2 weeks after the end of gavage.

### Cell Culture

HEK293T cells (ATCC, CRL-11268) were cultured in high-glucose Dulbecco’s Modified Eagle Medium (DMEM) supplemented with 10% FBS (Omega Scientific international, #881192) and antibiotics (HyClone, #SV30079.01). AML12 cells (ATCC, #CRL-2254) were cultured in DMEM: F12 (1:1) medium supplemented with 10% FBS, 40 ng/mL Dexamethasone (Gibco, A13449) and of 100x insulin-transferrin-selenium mixture (Gibco, #41400-045). Primary hepatocytes were isolated according to a two-step collagenase perfusion as follows: The 24G BD Nexia catheter (BD Biosciences, 383512) was inserted into the portal vein and the mouse was perfused with Ca²⁺/Mg²⁺-free HBSS (Hanks’ Balanced Salt Solution) (Gibco, 14175) containing 0.5 mM EGTA (Corning, 46034), then switched to 37 °C buffer of HBSS with Ca²⁺/Mg²⁺ (Gibco, 14025) with 5mM CaCl_2_ containing 0.1% (w/v) collagenase type IV. After perfusion, liver was gently teased apart and filtered into a tube containing cold DMEM with 70 µm cell strainer (Falcon, 352350). Cells were washed with cold DMEM 3 times and pelleted at 50 × g for 5 minutes at 4 °C. Hepatocytes were plated on collagen (Corning, #354236) -coated plates in warm high-glucose DMEM supplemented with 10% FBS and antibiotics. Primary LSECs were isolated using enzymes and were specifically enriched by MACS. Livers from 8-week-old male C57BL/6J mice were digested using a the gentleMACS Octo Dissociator (Miltenyi Biotech, 130-134-029) and Liver Dissociation Kit (Miltenyi Biotech, 130-105-807). CD45 positive cells were removed with CD45+ magnetic bead (Miltenyi Biotech, 130-052-301) LSECs were enriched by CD146+ magnetic bead selection (Miltenyi Biotech, 130-092-007), according to the manufacturer’s protocols. The cells were seeded on 0.1% gelatin (Sigma, #9000-70-8) coated plates in EGM-2 medium (EBM-2, Lonza, #CC-3156 with SingleQuots kit #CC-4176). All cells mentioned above were cultured in a humidified 37 °C, 5% CO₂ incubator. For further treatments, cells were washed with PBS for three times and starved in 0.2% FBS containing EBM-2 buffer for 6 hours. Later, the proteins and drugs as indicated were added to the cells in EBM-2 buffer containing 5% FBS.

### Plasmid construction and AAV production

To generate pAAV-TBG-mLIF expression vectors, the nucleic acid fragments encoding mouse LIF (Gene ID:16878) were synthesized by GenScript USA, Inc. and inserted with NotI/HindIII sites into pAAV.TBG.PI.eGFP.WPRE.bGH (Addgene #105535). For pAAV-CAG-Cre construct, Cre sequences were synthesized by GenScript USA, Inc. and then inserted into pAAV-CAG-GFP backbone (Addgene #37825) using GenBuilder TM Cloning Kit (GenScript, L00701). GFP backbones served as control for AAV vectors. The constructs were verified by sequencing and large-scale plasmid preparations were performed using plasmid maxiprep kit (Invitrogen, K210017). Helper plasmids (ΔF6, Addgene #112867; DJ, Cell Biolabs, VPK 400-DJ) and capsid plasmids (2/8, Addgene #112864; DJ, Cell Biolabs, VPK 400-DJ) were also amplified using the same kit. Plasmid quality was verified by agarose gel electrophoresis. AAV plasmids were transfected with PEI (ratio 1:4, DNA: PEI) into HEK293T cells for 60 h. Cells were harvested and resuspended in buffer (150 mM NaCl and 20 mM Tris pH8.0) and subjected to three freeze thaw cycles (liquid nitrogen/37 °C water bath) to release virus with 1 mM MgCl_2_ and 250 U/mL benzonase (Sigma, E8263). After centrifugation at 4000 rpm, 4 °C for 15 min, the supernatants were subjected to ultracentrifugation (Beckman, Optimal^TM^L-90K) at 53,000 rpm in 33 mL OptiSeal tubes (Beckman Coulter, 361625) with a discontinuous OptiPrep gradient (17-25-40-60%, Millipore, D1556). Viral particles were collected from the 40% layer and concentrated using 100 kDa Amicon® Ultra Centrifugal Filter (Millipore, UFC9100) in PBS containing 0.01% poloxamer188 (Corning, 13-901-CI). For the purity analysis, viral proteins were added 2× loading buffer (ThermoFisher, #B0007) with reducing agent (ThermoFisher, #B0009), and incubated at 95 °C for 5 min. The samples were analyzed by SDS-PAGE and silver-stained (Invitrogen, #45-1001) according to manufacturer’s protocol. Viral titers were quantified by qPCR using GFP plasmids (CAG or TBG backbones) with serial 10x dilution as standards. The qPCR was performed on a ViiA 7 relative standard curve system (Applied Biosystems) with 2 µL template, 10 µL 2× SYBR Green Master Mix (Vazyme, Q712-00), 0.5 µL forward/reverse primers and 7 µL ddH_2_O. Primer’s sequence: TBG Forward: TTCTGAGAGACTGCAGAAGTTG, Reverse: CTCGACAAGCCCAGTTTCTAT; CAG Forward: AGCCTCTGCTAACCATGTTC, Reverse: AAATGATGAGACAGCACAATAACC.

### AAV delivery

For in vivo LIF overexpression experiments, AAV-TBG-mLIF or GFP controls were delivered to 8-week-old male C57BL/6J mice through tail vein injection at indicated titers (diluted in 150 µL PBS). 30 days later, the mice were subjected to partial hepatectomy and sacrificed at different time points. The weights of spleen, kidney and liver were collected for the calculation of ratio normalized to body weight. The increased ratio after PHx was further calculated with the normalization to no surgery groups at indicated titers. For in vitro experiments, 5×10^11 GC virus of DJ-CAG-Cre was treated to the freshly isolated LSECs from *Lifr f/f* mice with the density 4×10^5 cells/mL for 48 hours, then the cells were ready for further treatments.

### Flow Cytometry

The freshly isolated LSECs were resuspended in FACS buffer (PBS with 2% FBS and 1 mM EDTA), containing TruStain FcX at the dilution of 1:500 (Biolegend,101319) and 1uM Zombie Violet Viability Dye (Biolegend, 423113), and then incubated at 4℃ in the dark for 10 minutes. After centrifugation at 300 g for 10min, the cells were resuspended in cocktail antibody buffer, including PE anti-mouse CD31 (Biolegend, 102507), FITC anti-mouse CD13 (Biolegend, 164009), BV711 anti-mouse CD45 (Biolegend, 103147) at the dilution of 1:50 in the dark for 20 minutes at 4℃. After centrifugation at 300 g for 10 minutes, cells were resuspended in 300 µL FACS buffer for FACS analysis (BD FACSymphony^TM^ A1). Single antibody was incubated with UltraComp eBeads Compensation Beads (Invitrogen,01-2222-42) for single-color compensation controls. The data was analyzed using FlowJo V10.

### Protein purification

D25 1.4 hybridoma cells (ATCC, #HB-11074) were cultured (Hybridoma-SFM, ThermoFisher Scientific, #12045084) to produce the anti-LIF antibody in large quantities. 1-2 L of cell culture supernatant was used for each batch of antibody purification. Conditioned media were centrifuged at 4000 g for 15 min and filtered through a 0.45 µm filter. The clarified supernatant was loaded on to the column (HiTrap™MabSelect™PrismA 5 mL, Cytiva) equilibrated in PBS. The column was thoroughly washed with the binding buffer, and then with binding buffer containing 0.5 M NaCl. The bound antibody was then eluted in 0.1 M citric acid, pH 3.2, and immediately neutralized by adding 1/5 th volume of 1 M Tris, pH 9.5. Antibody containing fractions were pooled, buffer exchanged in 20 mM Histidine acetate, pH 5.5, 150 mM NaCl using a desalting column HiPrep™26/10 desalting (GE Healthcare). Samples were concentrated by Amicon® Ultra-15 centrifugal filter Ultracel®-10k. Chromatography was performed in AKTA avant FPLC system (GE Healthcare). Instrument and column were sanitized by 0.5 N NaOH before each run as described (58). The purified antibody was quantified by ELISA (Mouse IgG antigen ELISA Kit, Molecular Innovations #MSIGGKT) and BCA protein estimation. The quality of the protein was determined by SDS-PAGE and silver staining. Endotoxin was measured by ToxiSensor chromogenic LAL Endotoxin assay kit (GenScript, #L00350) and was less than 0.05 EU/mg of protein.

### Antibody treatments

Mouse MAb IgG2a isotype control (BioXcell, C1.18.4, #BE0085) was diluted in PBS at the concentrations of 10 mg/kg. aLIF (purified at UCSD or from Genentech) was diluted in PBS at the concentrations of 20 mg/kg for intraperitoneal injection to 8-week-old male C57BL/6J mice. The antibodies were injected according to body weight every other day for twice, the last shot was 24 hours before PHx.

### Partial hepatectomy

70% PHx surgeries were carried out on mice around 8-10 week-old according to previous reference (59). In general, a midline abdominal skin and muscle incision about 3 cm long was made to expose the xiphoid process. The peritoneal cavity was retracted using retractors, thereby exposing the liver. Using a moistened cotton tip, the median was lifted and the left lateral lobe was held against the diaphragm/thorax. The silk thread was placed at the base of the left lateral lobe. Tying two ends of the suture over the top of the left lateral lobe and then the lobe was cut. The threads for the second knot were placed between the stump and the median lobe. The median lobe was pulled down over the suture. The tied median lobe above the suture was cut. After the resection of lobes, the peritoneum was closed with absorbable sutures (Ethicon, J213H). The peritoneum and skin were closed using the 9 mm Autoclips (Braintree Scientific, 205016). The mice were sacrificed at 24, 48 72 hours and 1 week after PHx.

### Proliferation assays

Freshly isolated LSECs were seeded at 7,000 cells/well in 0.1% gelatin-coated 96-well plates in low-glucose DMEM with 2% bovine calf serum, 2 mM glutamine, and antibiotics. Freshly isolated hepatocytes were seeded at 5,000 cells/well on 96-well plates in high-glucose DMEM containing 10% FBS with antibiotics. Before treatment, the primary hepatocytes were pre-incubated in high-glucose DMEM containing 0.5% FBS for 24 hours. Cells were than treated with DMEM: F12 (1:1) medium containing dexamethasone, insulin-transferrin-selenium and antibiotics. AML12 cells were seeded at 5,000 cells/well in 96-well plates in DMEM: F12 (1:1) medium with dexamethasone, insulin-transferrin-selenium and antibiotics. Before treatment, the cells were starved in DMEM: F12 (1:1) medium with 0.2% FBS for 24 hours, hLIF and HGF were diluted in the starving medium. The proteins which included hLIF (Abcam, #ab287941), hVEGF (R&D system, #293-VE), hOSM (R&D system, #8475-OM), hCT1 (R&D system, 612-CD-010), IL-6 (PeproTech, 200-06), hHGF(R&D system, 294-HG), mHGF (R&D system, 2207-HG-025), EGF (R&D system, 236-EG), aLIF (UCSD), MAb mouse IgG2a isotype control (BioXcell,C1.18.4, #BE0085), anti-VEGF antibody B20-4.1.1 (Genentech) were added to cells at various concentrations, as indicated in the figures. After 5 or 6 days, cells were incubated with 18 µL Resazurin solution (Biotium, 30025-2) for 4 h. Fluorescence was measured at 530-nm excitation wavelength and 590-nm emission wavelength using the plate reader SpectraMax M5 (Molecular Devices, Sunnyvale, CA).

### ELISA and liver enzyme activity assays

Blood was taken from mice in different experimental groups and collected in serum-separating tubes (BD, 365967), After clotting at room temperature for 1 hour, the serum was collected after centrifugation at 6000 rpm for 5 minutes at room temperature. For mLIF detection, the serum was pre-treated with 2.5 N acetic acid and then neutralized with 2.7 N NaOH/1 M HEPES. The following steps were carried out according to the manufacturer’s protocols from mLIF ELISA kit (R&D system, MLF00). For HGF detection, the serum was collected as above described and added to mHGF ELISA kit (Sigma-Aldrich, RAB0214) after dilution. The medium collected from freshly isolated LSECs with the treatment for 48 hours were centrifuged at 12,000 rpm for 15 minutes in Protein Lobind tubes (Sigma, EP022431081). The supernatant was applied directly to mHGF ELISA kit (R&D system, MHG00) according to the manufacturer’s protocols. For liver enzyme activity detection, the serum was collected as before and applied to AST (Cell Biolabs, MET-512t) and ALT (Cell Biolabs, MET-5123) kits after dilution.

### RNA isolation, reverse transcription and quantitative PCR

Total RNA was extracted using RNeasy Plus Micro Kit (Qiagen, 74034) or Trizol (Life Technologies, 15596018). Reverse transcription of equal amounts of RNA was performed with High-Capacity cDNA Reverse Transcription Kit (Applied Biosystems, 4368813), followed by quantitative PCR, using TaqMAn® Fast Advanced Master Mix (Applied Biosystems, 4444554) on a ViiA 7 real time PCR machine (Applied Biosciences, USA). mRNA levels were calculated using the ΔCt method and normalized to the internal control Actin. TaqMAn® expression assays were used: *Actb* (Mm02619580_g1), *Anpep* (Mm00476227_m1), *Cdh5* (Mm00486938_m1), *Hgf* (Mm01135184_m1), Kdr (Mm01222421_m1), *Lifr* (Mm00442942_m1), *Lyve1* (Mm00475056_m1), *Pdgfrb*, (Mm00435546_m1), Pecam (Mm01242576_m1).

### Immunoblotting

Isolated LSECs or hepatocytes were cultured in pre-coated plates for 24 hours. LSECs were then starved for 6 hours, while hepatocytes and Huvec cells were serum-starved for 24 hours, before addition of hLIF (40 ng/mL to LSECs, 100 ng/mL to hepatocytes) or hVEGF (50 ng/mL to LSECs and Huvec). After 10 minutes incubation at 37 °C, cells were harvested with RIPA lysis buffer (Thermo Scientific, 89901) containing protease and phosphatase inhibitors (Cell Signaling, 5872). The sample was boiled at 95 °C for 10 minutes with 4x Bolt LDS Sample Buffer (Life Technologies, B0007) and 10 x Sample Reducing Agent (Life Technologies, B0009), then loaded to 4-12% SDS-PAGE gels (Life Technologies, NW04125BOX). For LIFR detection, the samples were loaded to the 3-8% Tris-acetate gels (Invitrogen, EA03785). The gels were transferred to PVDF membrane (Millipore, IPFL00010) and were blocked with blocking buffer (Licor, 92760001) and incubated with primary antibody at 4 °C overnight, followed by incubation with the secondary antibody conjugated with fluorescence, goat anti mouse IRDye 800CW and goat anti rabbit IRDye680RD, (Licor, 926-32210 and 926-32211). The antibodies were diluted in the dilution buffer (Licor, 927-65001). The images were captured by a Licor Odyssey imaging system (Licor, USA). The following primary antibodies were used in this study : LIFR (Proteintech, 22779-1-AP, 1:1,000 dilution), Tubulin (Proteintech, 66031-1-Ig, 1:2,000 dilution), Actin (Proteintech, 66009-1-Ig, 1:10,000 dilution), PCNA (Abcam, AB220208, 1:2,000 dilution), p-Y1175-VEGFR2 (Cell Signaling Technology, 3770S, 1:2000 dilution), Total-VEGFR2 (Cell Signaling Technology, 2479S, 1:2000 dilution), p-T705-STAT3 (Cell Signaling Technology, 9145S, 1:2000 dilution), Total-STAT3 (Cell Signaling Technology, 30835S, 1:2,000 dilution), p-T202/Y204-ERK (Cell Signaling Technology, 4370, 1:2,000 dilution) and Total-ERK (Cell Signaling Technology, 4695, 1:2,000 dilution), p-S473-AKT (Cell Signaling Technology, 4060, 1:2,000 dilution), Total-AKT (Cell Signaling Technology, 2920, 1:2000 dilution)

### Co-culture of LSECs and hepatocytes

Isolated hepatocytes were seeded into 24-well plate at the density of 130,000 cells/ well in high-glucose DMEM containing 10% FBS and antibiotics. The freshly isolated LSECs were seeded on the Transwell inserts (Costar, 3422) at the density of 50,000 cells/well separately with EGM-2 medium. The primary hepatocytes were serum-starved for 24 hours in high-glucose DMEM containing 0.5% FBS. The LSECs were starved for 6 hours in EBM-2 containing 0.2% FBS, then transferred to 200 µL EBM2 buffer with 5% FBS and co-cultured with hepatocytes seeded in 24-well plate with 400 µL DMEM/F12 medium containing with 5% FBS, dexamethasone, insulin-transferrin-selenium and antibiotics. The co-culture plates were treated with 80 ng/mL LIF or not for 4 days. Then the medium was collected for ELISA detection after centrifuge. The hepatocytes were harvested by same volume of RIPA containing protease and phosphatase inhibitors for immunoblot.

### Histology and immunofluorescence staining

Mice were euthanized and the livers were removed and fixed in 4% paraformaldehyde (Thermo scientific, J19943-K2) for 48 hours, transferred to 70% ethanol and embedded in paraffin, sectioned and stained with hematoxylin and eosin (H&E) by the Histology Core of Biorepository and Tissue Technology Shared Resources (TTSR) at Moores Cancer Center at UCSD. 5 µm thick sections were used for immunofluorescence staining, which were deparaffinized in xylene and an ethanol series, followed by two 5 minutes rinses in PBS. Antigen retrieval was performed with citrate buffer (Millipore Sigma, C9999) in a pressure cooker for 4 minutes and cool down to room temperature before two more 5-minute rinses in PBS. Sections were then circled with a hydrophobic barrier pen and blocked with 10% goat serum in PBS for 1 h. For AML12 immunofluorescence staining, the cells were plated on the 35 mm glass dishes, after treatment of HGF or LIF for 48 hours, the cells were fixed in methanol for 5 minutes, permeabilized with 0.1% Triton X-100 for 5 minutes and then blocked with 1% BSA/10% normal goat serum/0.3 M glycine in 0.1% PBS-Tween20 for 1 h. For all stainings, incubation with primary antibody was done overnight at 4℃ in a humidified chamber. Samples were stained with rabbit anti cytokeratin 19 (Abcam, AB133496, 1:100 dilution) and PCNA (Abcam, AB220208, 1:50). AML12 samples were stained with Ki67 (Abcam, AB16667, 1:25 dilution). After washing with PBS fluorescent secondary antibodies were applied in a dilution of 1:500 (all from Invitrogen, Goat anti-Rabbit IgG (H+L) Cross-Adsorbed Secondary Antibody, Alexa Fluor™ 488, A-11008; Goat anti-Mouse IgG (H+L) Highly Cross-Adsorbed Secondary Antibody, Alexa Fluor™ 555, A-21424). After final washes with PBS slides were mounted with DAPI containing Fluoroshield mounting medium (Millipore Sigma, F6057) and imaged with AT2 Aperio slide scanner (Leica, Germany) or Keyence Microscope BZ-X710 (Keyence Corporation, Osaka, Japan). AML12 samples were imaged with confocal microscope (NIKON A1R confocal TIRF STORM microscopy). Analysis of acquired pictures was done in ImageJ using the cell counter plugin.

### Sample preparation for secretome analysis

Freshly isolated LSECs were seeded at 4×10^5 cells/well on 0.1% gelatin-coated 12-well plates in EGM-2 buffer. After a 6-hour starvation in EBM-2 buffer containing 0.2% FBS, the cells were treated with 40 ng/mL LIF in EBM-2 buffer supplemented with 5% FBS for 48 hours. Media were collected in Protein Lobind tubes (Sigma, EP022431081) and centrifuged at 12,000 rpm for 15 minutes. Samples were processed with the Proteograph XT Assay (60) (61) in Seer Inc. for later analysis. In brief, 240 µL medium were transferred to Seer Sample Tubes for processing with the Proteograph XT Assay kit (S55R1100). Proteins were quantitatively captured in nanoparticle (NP) associated protein coronas and subsequently denatured, reduced, alkylated and subjected to proteolytic digestion (trypsin and Lysc). Peptides were purified and yields were determined (Thermo Fisher Scientific catalog #23290). Peptides were dried down overnight with a vacuum concentrator and reconstituted with a reconstitution buffer to a concentration of 50 ng/µL.

### Data-Independent Acquisition LC-MS/MS

For Data-Independent Acquisition (DIA), 8 µL of reconstituted peptide mixture from each NP preparation was analyzed resulting in a constant 400 ng mass MS injection between NP A and NP B samples. Each sample was analyzed with a Vanquish NEO nanoLC system coupled with a Orbitrap TM Astral TM (Thermo Fisher, Germany) mass spectrometer using a trap-and-elute configuration. First, the peptides were loaded onto an AcclaimTM PepMapTM 100 C18 (0.3 mm ID x 5 mm) trap column and then separated on a 50 cm µPACTM analytical column (PharmaFluidics, Belgium) at a flow rate of 1 µL/min using a gradient of 5 - 25%solvent B (0.1% FA, 100% ACN) mixed into solvent A (0.1% FA, 100% water) over 20 minutes, resulting in a 24 minutes total run time. The mass spectrometer was operated in DIA mode with MS1 scanning and MS2 precursor isolation windows between 380-980 m/z. MS1 scans were performed in the Orbitrap detector at 240,000 R every 0.6 seconds with a 5 ms ion injection time or 500% AGC (500,000 ion) target. Two-hundred fixed window MS2 DIA scans were collected at the Astral detector per cycle with 3 Th precursor isolation windows, 25% normalized collision energy, and 5 ms ion injection times with a 500% (50,000 ion) active gain control maximum. MS2 scans were collected from 150-2000 m/z.

### DIA Raw Data Processing

DIA data was processed using ProteographTM Analysis Suite. Raw MS data was processed using the DIA-NN search engine (version 1.8.1) in library-free mode searching MS/MS spectra against an in silico generated spectral library of human protein entries (UP000005640_9606). Library-free search parameters include trypsin protease, 1 missed cleavage, N-terminal Met excision, fixed modification of Cys carbamidomethylation, no Met oxidation, peptide length of 7-30 amino acids, precursor range of 300-1800 m/z, and fragment ion range of 200-1800 m/z. MS1 and MS2 mass accuracy was set to 10 ppm. Precursor and Protein Group FDR thresholds were set at 1%. Quantification was performed on summed abundances of all unique peptides considering only precursors passing the FDR thresholds. PAS summarizes all nanoparticle values for a single protein into a single quantitative value. Specifically, a single protein may have been measured up to five times, once for each nanoparticle. To derive the single measurement value, PAS uses a maximum representation approach, whereby the single quantification value for a particular peptide or protein group represents the quantitation value of the NP most frequently measured across all samples.

### Transcriptomics analysis

All human snRNA-seq analyses were performed in Python (version 3.10) using Scanpy (v1.10), numpy (v1.26), pandas (v2.2), matplotlib (v3.8), and seaborn (v0.13) for preprocessing, visualization, and statistical analysis. Raw .h5ad files from each sample were loaded using Scanpy, and all samples were subset to a shared gene set to ensure a harmonized feature space before merging. Batch correction was performed using scVI (scvi-tools v1.1.2). For each group (Healthy and APAP), we trained an scVI model with the following parameters: n_layers = 2, n_latent = 20, max_epochs = 50, batch_key = “batch”. To aggregate expression at the cell-type level, scVI-corrected gene expression values were averaged across all cells within each annotated cell-type category. These aggregated values were used for downstream ligand-receptor analysis. Cell-cell communication was quantified using a simple interaction model. Directional interaction scores were computed for all sender-receiver cell-type combinations by multiplying the mean expression of the ligand in the sending cell type by the mean expression of the corresponding receptor in the receiving cell type. Circular interactome visualizations were generated using networkx (v3.2) for directed graph construction, matplotlib for node and edge rendering, and dynamic percentile-based edge filtering was applied using a 90th percentile cutoff. Node size was proportional to cell counts per cell type, and node color represented ligand-receiver expression bias. Scatter plots were generated using matplotlib and seaborn. Sankey diagrams were generated with plotly (v5.20). Bubble plots were generated with holoviews (v1.17)

Human spatial transcriptomics (10x Visium) CSV files (spot metadata and gene expression) were processed from Seurat objects in R, and Python was subsequently used with pandas, numpy, and Scanpy (v1.10) to generate all the required visualizations. Seurat-provided spatial clusters were used directly without re-clustering. For each spatial Seurat cluster, marker genes were identified using a non-parametric Wilcoxon rank-sum test comparing each cluster vs. all other cells and the top 20 genes per cluster (lowest p-values) were retained for downstream inference of biological domain identities.

Mouse datasets were processed identically to the human snRNA workflows except for batch correction. Data was loaded using Scanpy and no batch correction was applied because samples were analyzed individually. In Fig. S1 A, mean log-normalized ligand expression is shown for NC1 and CCL_4_ conditions, with a logarithmic y-axis used to enhance visualization across a broad dynamic range. Cell types were assigned using CellTypist (v1.5) with the Healthy_Mouse_Liver pretrained model “Healthy_Mouse_Liver.pkl”.

## Statistics and reproducibility

Data were presented as mean ± SEM. Proliferation data were presented as mean ± SD. Statistical significance was determined using unpaired, two-tailed Student’s t-tests. P values <0.05 were considered significant. All quantification were analyzed in ImageJ (FiJi). All graph figures were generated in GraphPad Prism software. R was used for the differential analysis of secretome data. Python was used for transcriptomics analysis of human and mouse liver samples. Except for animal studies (one or two times) and secretome (two times), each experiment was repeated at least three times with similar results.

## Acknowledgments

We thank Dr.Kyunghee Choi for providing *Lifr f/f* mice, generated by Dr. Colin Stewart. We thank Dr. Miguel Lopez-Ramirez for providing VE-Cadherin-Cre mice which were generated by Dr. Ralf H. Adams. We thank the UCSD Animal Care Facilities for excellent support. We are grateful to Dr. Erpei Wang from UCSD Moores Cancer Center Shared Resources for her professional help on Bioinformatics analysis. We thank Genentech, Inc. for the gift of anti-LIF antibody D.25 and anti-VEGF antibody B.20.4.1. We thank the Moores Cancer Center Microscopy Facility, Flow Cytometry Shared Resources and Histology Core center for their excellent support. The graphic abstract was created in https://BioRender.com. We thank NIH funding grant R35GM149261, the content of this paper does not necessarily represent the official views of the NIH.

