## Supplemental figures for "The LIF-LIFR Axis Promotes Liver Regeneration via Modulation of Angiogenesis and HGF Release from LSECs"

**Supplementary figure legends**

**Figure S1.**

(A) ligands expression in the scRNA-seq from mouse livers with carbon tetrachloride ($CCl_4$)-induced injury.

(B) *vegfa* expression in the scRNA-seq across cell types in the livers of mouse with APAP injury at the indicated time points.

(C) *LIF* expression in the scRNA-seq from human livers with APAP injury.

(D&E) *LIF* (D) and *VEGFA* (E) expression in the scRNA-seq across cell types in the human liver samples with APAP injury.

(F) *LIF* expression in the spatial transcriptomics of liver samples from healthy (HEA2) and APAP injury (APAP1_S2) humans.

(G) Resected liver-to-body weight ratio in male mice treated with anti-LIF antibody (Genentech) or control IgG.

(H) Liver-to-body weight ratio at 24 h after partial hepatectomy (PHx) in male mice treated with anti-LIF antibody (Genentech) or control IgG.

(I) H&E staining of liver sections from mice treated with anti-LIF antibody or control IgG. Scale bars: 300 μm.

(J) Serum AST levels at 24, 48, 72 h and 1-week post-PHx in mice treated with anti-LIF antibody or control IgG.

**Figure S2.**

(A) *lifr* expression in scRNA-seq across cell types of liver from healthy mice (Adult control).

(B) Expression of IL-6 family receptors in scRNA-seq across cell types from healthy human liver samples. Upper is bar plot of *LIFR* expression, lower is bubble plot of multiple receptors as indicated.

(C) Circular network plot of *LIF-LIFR* interaction between different cell types in liver from healthy human (HEA8).

(D&E) Sankey plots of *LIF-LIFR* interaction between different cell types in liver from healthy human (HEA8 and HEA5).

(F) qPCR of LSEC markers (*Vegfr2, Cd31, Cdh5*) and markers of non-endothelial populations (including *Anpep, Pdgfrb, Lyve1*) in freshly isolated primary LSECs.

(G) Flow cytometry analysis of freshly isolated LSECs using CD31-PE antibody, CD13-FITC, CD45-BV711.

(H) qPCR of *lifr* in whole liver (Left) and isolated primary LSECs (Right) at indicated time points post-PHx.

**Figure S3.**

(A) Proliferation assays of LSECs treated with increasing amounts of IL-6; 100 ng/mL LIF was included as positive control.

(B-D) Proliferation assays of LSECs treated with increasing concentrations of anti-LIF antibody (B) or control IgG (D) in the presence of 10 ng/mL hLIF, and assessment of cross-inhibition on 10 ng/mL hVEGF (C).

(E&F) Proliferation assays of LSECs treated with increasing of anti-VEGF antibody B.20.4.1 in the presence of 10 ng/mL hVEGF (E) or 10 ng/mL hLIF (F).

(G) Proliferation assays of primary hepatocytes treated with increasing concentrations of hHGF and hEGF.

(H) Proliferation assays of AML12 cells treated with increasing concentrations of hHGF.

**Figure S4.**

(A) PCA analysis of LSECs secretome treated with 40 ng/mL hLIF versus control (n = 4).

(B) qPCR of *hgf* in LSECs treated with hLIF, hCT-1 or hOSM isolated from *lifr f/f* or *lifr*Δ*EC* mice.

(C) Western blot quantification of hepatocytes co-cultured with LSECs with or without 40 ng/mL LIF.

(D&E) Proliferation assays of LSECs treated with increasing concentrations of mHGF (D) or hHGF (E).

**Figure S5.**

(A&B) Spleen weight (A) and spleen-to-body weight ratio (B) of mice injected with AAV8-TBG-mLIF at the indicated titers.

(C&D) Kidney weight (C) and kidney-to-body weight ratio (D) of mice injected with AAV8-TBG-mLIF at the indicated titers.

(E) H&E stained liver sections from mice injected with low-doses AAV8-TBG-mLIF ($1\times10^8$ or $2.5\times10^8$ GC/mouse).

(F) Increased liver-to-body weight ratio post-PHx at the indicated time points compared to no surgery controls in mice injected with low-dose AAV8-TBG-mLIF ($1\times10^8$ or $2.5\times10^8$ GC/mouse).

(G) Serum AST levels after PHx at the indicated time points in mice injected with low-dose AAV8-TBG-mLIF ($1\times10^8$ or $2.5\times10^8$ GC/mouse).

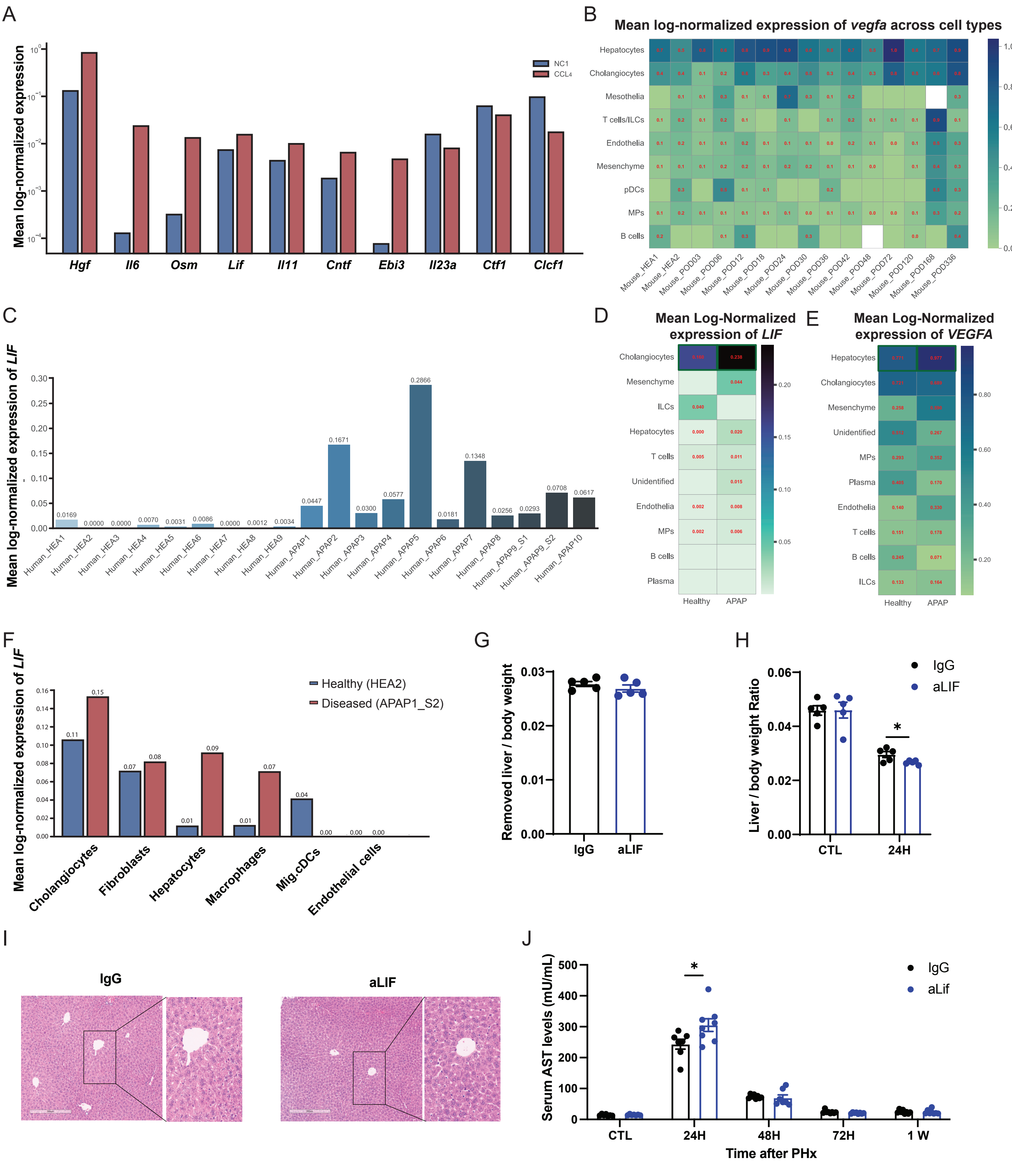
A
Mean log-normalized expression
NC1
CCL4
Hgf
Il6
Osm
Lif
Il11
Cntf
Ebi3
Il23a
Ctf1
Clcf1
B
Mean log-normalized expression of vegfa across cell types
Hepatocytes
Cholangiocytes
Mesothelia
T cells/ILCs
Endothelia
Mesenchyme
pDCs
MPs
B cells
Mouse_HEA1
Mouse_HEA2
Mouse_POD03
Mouse_POD06
Mouse_POD12
Mouse_POD18
Mouse_POD24
Mouse_POD30
Mouse_POD36
Mouse_POD42
Mouse_POD48
Mouse_POD72
Mouse_POD120
Mouse_POD168
Mouse_POD336
C
Mean log-normalized expression of LIF
0.0169
0.0000
0.0000
0.0070
0.0031
0.0086
0.0000
0.0012
0.0034
0.0447
0.1671
0.0300
0.0577
0.2866
0.0181
0.1348
0.0256
0.0293
0.0708
0.0617
Human_HEA1
Human_HEA2
Human_HEA3
Human_HEA4
Human_HEA5
Human_HEA6
Human_HEA7
Human_HEA8
Human_HEA9
Human_APAP1
Human_APAP2
Human_APAP3
Human_APAP4
Human_APAP5
Human_APAP6
Human_APAP7
Human_APAP8
Human_APAP9_S1
Human_APAP9_S2
Human_APAP10
D
Mean Log-Normalized expression of LIF
Cholangiocytes
Mesenchyme
ILCs
Hepatocytes
T cells
Unidentified
Endothelia
MPs
B cells
Plasma
Healthy
APAP
E
Mean Log-Normalized expression of VEGFA
Hepatocytes
Cholangiocytes
Mesenchyme
Unidentified
MPs
Plasma
Endothelia
T cells
B cells
ILCs
Healthy
APAP
F
Mean log-normalized expression of LIF
Healthy (HEA2)
Diseased (APAP1_S2)
0.11
0.15
0.07
0.08
0.01
0.09
0.01
0.07
0.04
0.00
0.00
0.00
Cholangiocytes
Fibroblasts
Hepatocytes
Macrophages
Mig.cDCs
Endothelial cells
G
Removed liver / body weight
IgG
aLIF
H
Liver / body weight Ratio
IgG
aLIF
*
CTL
24H
I
IgG
aLIF
J
Serum AST levels (mU/mL)
IgG
aLif
*
CTL
24H
48H
72H
1 W
Time after PHx

**Figure S1**

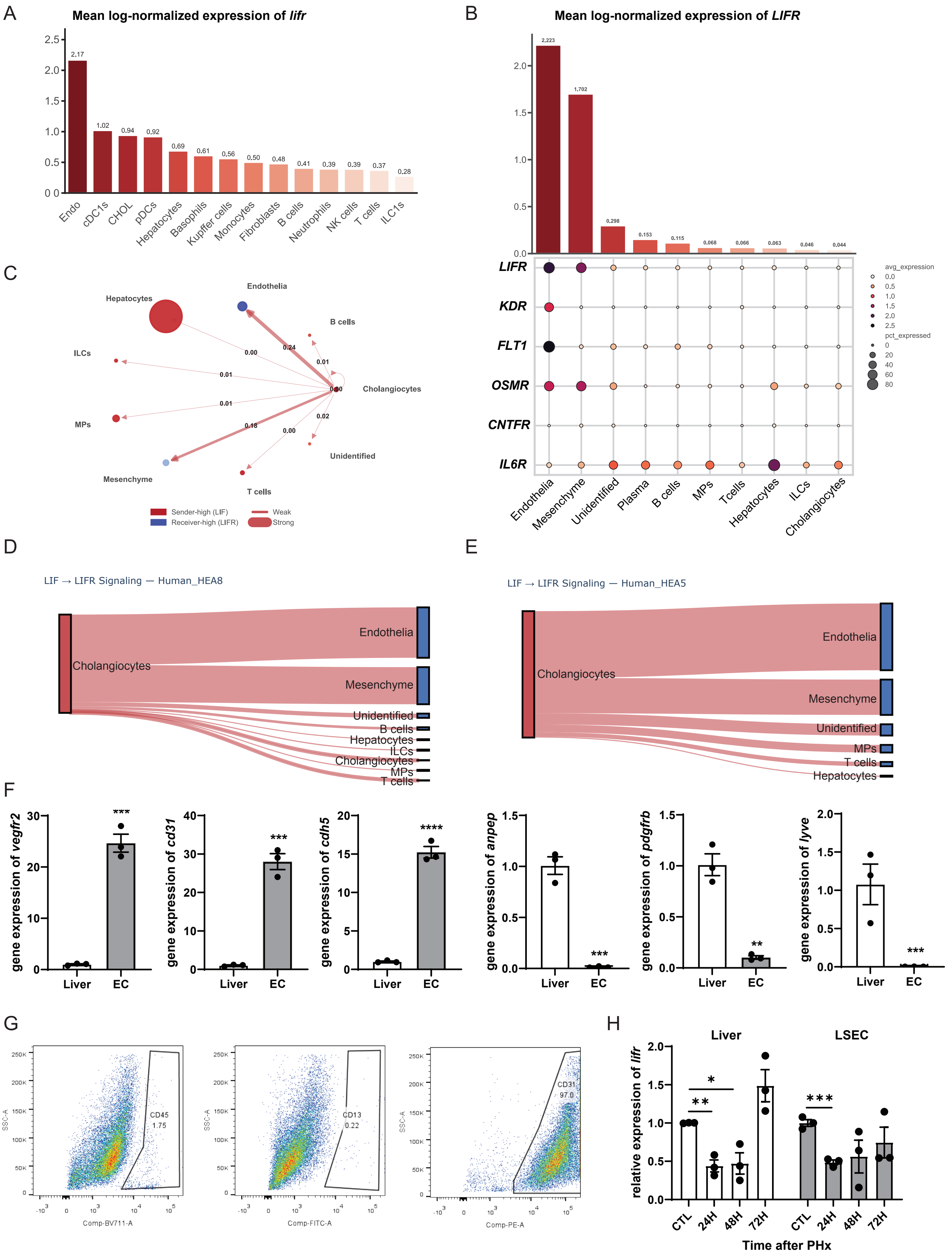
A
Mean log-normalized expression of lifr
2.17
1.02
0.94
0.92
0.69
0.61
0.56
0.50
0.48
0.41
0.39
0.39
0.37
0.28
Endo
cDC1s
CHOL
pDCs
Hepatocytes
Basophils
Kupffer cells
Monocytes
Fibroblasts
B cells
Neutrophils
NK cells
T cells
ILC1s
B
Mean log-normalized expression of LIFR
2.223
1.702
0.298
0.153
0.115
0.068
0.066
0.063
0.046
0.044
LIFR
KDR
FLT1
OSMR
CNTFR
IL6R
Endothelia
Mesenchyme
Unidentified
Plasma
B cells
MPs
Tcells
Hepatocytes
ILCs
Cholangiocytes
avg_expression
pct_expressed
C
Hepatocytes
Endothelia
B cells
ILCs
Cholangiocytes
MPs
Unidentified
Mesenchyme
T cells
Sender-high (LIF)
Receiver-high (LIFR)
Weak
Strong
D
LIF → LIFR Signaling — Human_HEA8
Cholangiocytes
Endothelia
Mesenchyme
Unidentified
B cells
Hepatocytes
ILCs
Cholangiocytes
MPs
T cells
E
LIF → LIFR Signaling — Human_HEA5
Cholangiocytes
Endothelia
Mesenchyme
Unidentified
MPs
T cells
Hepatocytes
F
gene expression of vegfr2
gene expression of cd31
gene expression of cdh5
gene expression of anpep
gene expression of pdgfrb
gene expression of lyve
Liver
EC
G
CD45
1.75
CD13
0.22
CD31
97.0
SSC-A
Comp-BV711-A
Comp-FITC-A
Comp-PE-A
H
Liver
LSEC
relative expression of lifr
CTL
24H
48H
72H
Time after PHx

Figure S2

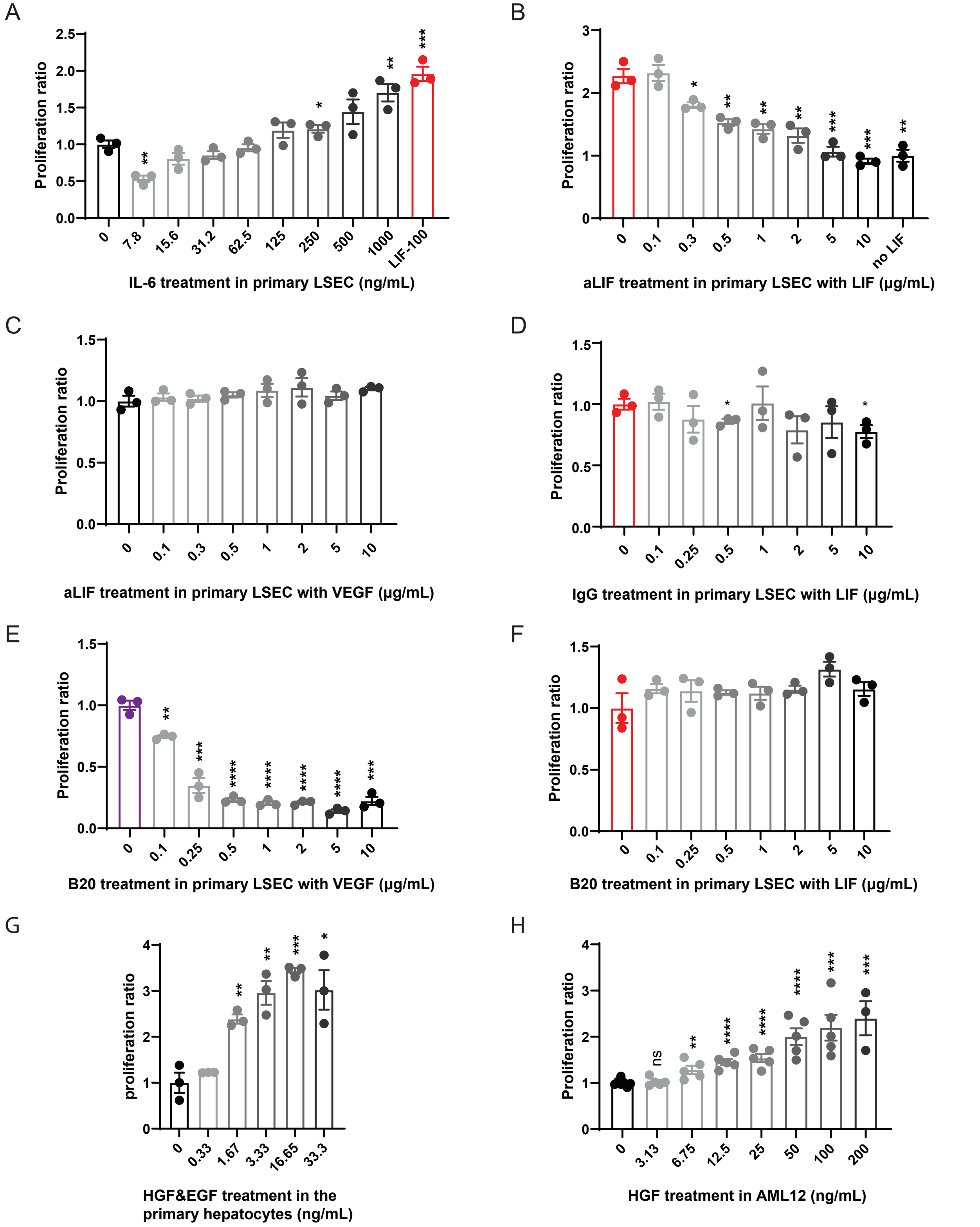
A
Proliferation ratio
2.5
2.0
1.5
1.0
0.5
0.0
0
7.8
15.6
31.2
62.5
125
250
500
1000
LIF-100
IL-6 treatment in primary LSEC (ng/mL)
B
Proliferation ratio
no LIF
aLIF treatment in primary LSEC with LIF (μg/mL)
C
Proliferation ratio
aLIF treatment in primary LSEC with VEGF (μg/mL)
D
Proliferation ratio
IgG treatment in primary LSEC with LIF (μg/mL)
E
Proliferation ratio
B20 treatment in primary LSEC with VEGF (μg/mL)
F
Proliferation ratio
B20 treatment in primary LSEC with LIF (μg/mL)
G
proliferation ratio
0
0.33
1.67
3.33
16.65
33.3
HGF&EGF treatment in the
primary hepatocytes (ng/mL)
H
Proliferation ratio
ns
0
3.13
6.75
12.5
25
50
100
200
HGF treatment in AML12 (ng/mL)

**Figure S3**

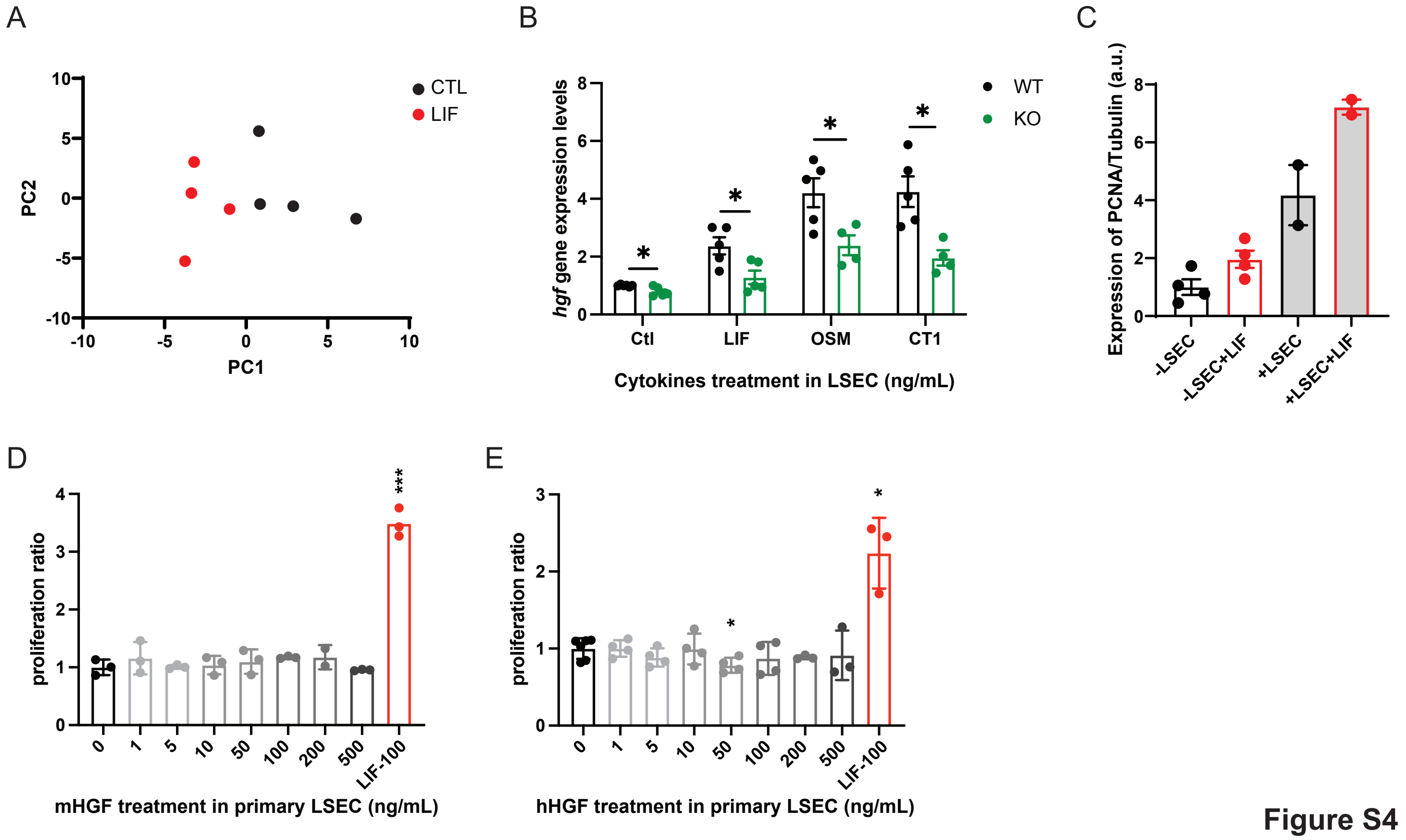
A
CTL
LIF
PC2
PC1
B
WT
KO
hgf gene expression levels
Ctl
LIF
OSM
CT1
Cytokines treatment in LSEC (ng/mL)
C
Expression of PCNA/Tubulin (a.u.)
-LSEC
-LSEC+LIF
+LSEC
+LSEC+LIF
D
proliferation ratio
0
1
5
10
50
100
200
500
LIF-100
mHGF treatment in primary LSEC (ng/mL)
E
proliferation ratio
hHGF treatment in primary LSEC (ng/mL)

Figure S4

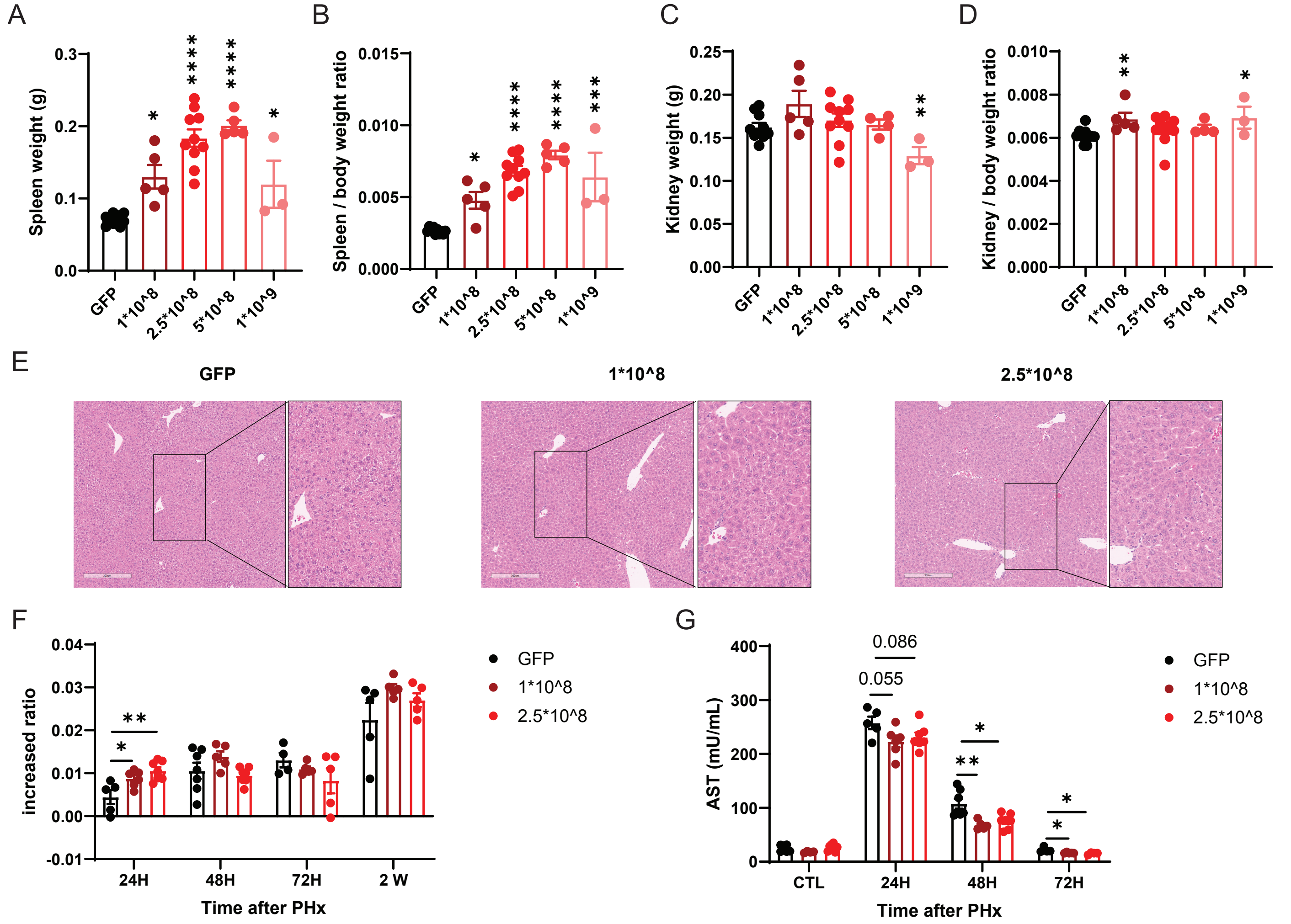
A
Spleen weight (g)
B
Spleen / body weight ratio
C
Kidney weight (g)
D
Kidney / body weight ratio
GFP
1*10^8
2.5*10^8
5*10^8
1*10^9
E
GFP
1*10^8
2.5*10^8
F
increased ratio
24H
48H
72H
2 W
Time after PHx
GFP
1*10^8
2.5*10^8
G
AST (mU/mL)
0.086
0.055
CTL
24H
48H
72H
Time after PHx
GFP
1*10^8
2.5*10^8

**Figure S5**
